# Sequential *in-cis* mutagenesis *in vivo* reveals various functions for CTCF sites at the mouse *HoxD* cluster

**DOI:** 10.1101/2021.08.13.456193

**Authors:** Ana Rita Amândio, Leonardo Beccari, Lucille Lopez-Delisle, Bénédicte Mascrez, Jozsef Zakany, Sandra Gitto, Denis Duboule

## Abstract

Mammalian *Hox* gene clusters contain a range of CTCF binding sites. In addition to their importance in organizing a TAD border, which isolates the most posterior genes from the rest of the cluster, the positions and orientations of these sites suggest that CTCF may be instrumental in the selection of various subsets of contiguous genes, which are targets of distinct remote enhancers located in the flanking regulatory landscapes. We examined this possibility by producing an allelic series of cumulative *in-cis* mutations in these sites, up to the abrogation of CTCF binding in the five sites located on one side of the TAD border. In the most impactful alleles, the global chromatin architecture of the locus was modified, yet not drastically, illustrating that CTCF sites located on one side of a strong TAD border are sufficient to organize at least part of this insulation. Spatial colinearity in the expression of these genes along the major body axis was nevertheless maintained, despite abnormal expression boundaries. In contrast, strong effects were scored in the selection of target genes responding to particular enhancers, leading to the mis-regulation of *Hoxd* genes in specific structures. Altogether, while most enhancer-promoter interactions can occur in the absence of this series of CTCF sites, it seems that the binding of CTCF in the *Hox* cluster is required to properly transform a rather unprecise process into a highly discriminative mechanism of interactions, which is translated into various patterns of transcription accompanied by the distinctive chromatin topology found at this locus. Our allelic series also allowed us to reveal the distinct functional contributions for CTCF sites within this *Hox* cluster, some acting as insulator elements, others being necessary to anchor or stabilize enhancer-promoter interactions and some doing both, whereas all together contribute to the formation of a TAD border. This variety of tasks may explain the amazing evolutionary conservation in the distribution of these sites amongst paralogous *Hox* clusters or between various vertebrates.

## INTRODUCTION

Embryonic development relies on complex and precise dynamics of gene activation and repression, driven in large part by the combined activity of multiple *cis*-regulatory elements (CREs) (Long et al., 2016; Spitz and Furlong, 2012). In vertebrates, CREs can be located at long distances from their target genes and interact with them through the establishment of particular chromatin structures such as loops. Often, the same genomic region harbors multiple regulatory elements and several transcription units, raising the question as to how specific enhancer-promoter interactions can be established without affecting neighboring genes.

The advent of chromosome conformation capture (3C) technologies (Dekker, 2006) confirmed that the eukaryote genome is organized into several levels of folding with, at the megabase level, chromatin domains referred to as topologically associating domains or TADs (Dixon et al., 2012; Nora et al., 2012; Sexton et al., 2012). TADs are domains where DNA sequences such as promoters and their enhancers interact more frequently than with regions located outside, independently from the linear distance (Dixon et al., 2016, 2012) and may thus constitute structural units in the organization of genomes associated with particular functional tasks. Indeed, complex regulatory landscapes spread over large distances often match TADs (e.g. (Andrey et al., 2013). Even though a causal relationship remains to be fully clarified, TADs are thus thought to delimit functionally autonomous regions, somewhat channeling the activity of distal CREs (Sikorska and Sexton, 2020) by reducing the search space between the enhancers and their promoters (Symmons et al., 2016). Accordingly, these domains tend to be evolutionary conserved within large syntenic regions (Dixon et al., 2012; Krefting et al., 2018). In support of this view, the disruption of TAD borders was shown to lead to their loss of insulation and concurrent mis-regulation of genes due ectopic gene-enhancer interactions (Gómez-Marín et al., 2015; Ibn-Salem et al., 2017; Lupianez et al., 2015; Rodriguez-Carballo et al., 2017). In agreement, genomic rearrangements whereby TAD boundaries are placed between regulatory elements and their targets genes result in the downregulation of the target genes along with TAD reorganization (Kraft et al., 2019; Lupianez et al., 2015; Willemin et al., 2021). However, how tissue- or gene-specific contacts can be established within one TAD is still elusive, in particular how distinct sets of enhancer sequences can interact with various subsets of transcription units and not others, in different cell types, while all located in the same chromatin domain.

Amongst those proteins that contribute to the establishment of the nuclear 3D chromatin organization, the CTCF zing-finger transcription factor plays an important role. It was initially described as a negative regulator of gene expression, due to its capacity to repress transcription by blocking enhancer-promoter interactions, thus defining a category of CREs referred to as insulators (Bell et al., 1999; Chung et al., 1993; Lobanenkov et al., 1990), see (Herold et al., 2012). CTCF recognizes a conserved GC-rich 20 nucleotides long consensus sequence (Nakahashi et al., 2013; Renda et al., 2007; Yin et al., 2017) and mediates loop formation in conjunction with Cohesin, a protein complex with chromatin extruding activity (Hansen et al., 2017a; Merkenschlager and Odom, 2013; Sedeño Cacciatore and Rowland, 2019). In the ‘loop-extrusion’ model, Cohesin is loaded into the chromatin where it forms a ring-shaped structure which moves along and progressively extrudes the DNA fiber until reaching CTCF-occupied sites where the CTCF N-terminal portion faces Cohesin progression (Davidson et al., 2019; Fudenberg et al., 2016; Hansen et al., 2017b; Kim et al., 2019; Pugacheva et al., 2020; Sanborn et al., 2015; Stigler et al., 2016; Xi and Beer, 2021).

Accordingly, the orientation and location of the CTCF binding sites (CBSs) play a critical role in DNA loop formation and high-order chromatin organization. In fact, TAD and sub-TADs boundaries are enriched in CBSs, with usually several CTCF motifs displaying the same orientation, facing those sites located at the other extremity of the TAD (Huang et al., 2021; Kentepozidou et al., 2020). Despite their prominent role in the establishment of TAD boundaries, most CBSs are found outside these regions and are associated with a wide range of functions including enhancer-promoter interaction, imprinting and recombination (Franco et al., 2014; Gosalia et al., 2014; Guo et al., 2011; Phillips-Cremins et al., 2013). In agreement with this multifaceted role, CTCF depletion in the embryo resulted in the concomitant loss of TAD insulation and weakening of genes-enhancers interactions, sometimes with clearly documented effects upon gene transcription (e.g. (Paliou et al., 2019) whereas in other instances a more moderate and somewhat unpredictable impact was observed (Luan et al., 2021; Nora et al., 2017; Soshnikova et al., 2010). However, the potential importance of CTCF in helping tissue specific enhancers to select the right promoter(s) and thus activate a subset of genes located within the same TAD remains to be assessed with precision. In this context, *Hox* gene clusters provide an excellent experimental paradigm. Indeed besides their critical function in the organization of the major body axis (Wellik, 2009), these genes are highly pleiotropic as they are involved in the making of a range of organs and structures at various developmental times (e.g.(Krumlauf, 1994). This is exemplified by the limbs, the external genitals, the uro-genital and gastro- intestinal tractus (Deschamps and Duboule, 2017; Favier and Dollé, 1997; Mallo et al., 2010; Zakany and Duboule, 2007), as well as other endodermal organ, or teguments such as hairs (Godwin and Capecchi, 1998) and nails (Fernandez-Guerrero et al., 2020). The various enhancers necessary to achieve these widely diverse regulations are positioned on either sides of the clusters and have been characterized in some details, in particular at the mouse *HoxD* locus.

This locus, which included nine genes within a ca 100kb large DNA segment, is positioned in between two large regulatory domains matching TADs (C-DOM and T-DOM) and contains in itself a strong chromatin boundary, a structure that leads to a differential tropism in enhancers-promoters interactions (Andrey et al., 2013; Darbellay and Duboule, 2016; Noordermeer et al., 2011; Rodriguez-Carballo et al., 2017); Fig. 1A). While the ‘posterior’ genes *Hoxd13* and *Hoxd12* mainly contact the C-DOM, the ‘anterior’ part of the cluster (from *Hoxd1 to Hoxd9*) preferentially interacts with the T-DOM, with *Hoxd10* and *Hoxd11* being more versatile in their interaction potential. This internal chromatin boundary is induced by the presence of a collection of nine CTCF sites, with an inversion of polarity in the middle, which positions the terminal genes *Hoxd12* and *Hoxd13* in a TAD (C-DOM) that is distinct from that containing the rest of the cluster (T-DOM; Fig. 1C). While C-DOM contains several enhancers necessary to produce ‘terminal’ structures, the hands and feet, as well as the genitals (Amândio et al., 2019; Montavon et al., 2011), T-DOM includes range of enhancer sequences specific for various structures such as the proximal limbs, part of the intestines, mammary glands or various head structures.

**Figure 1:**
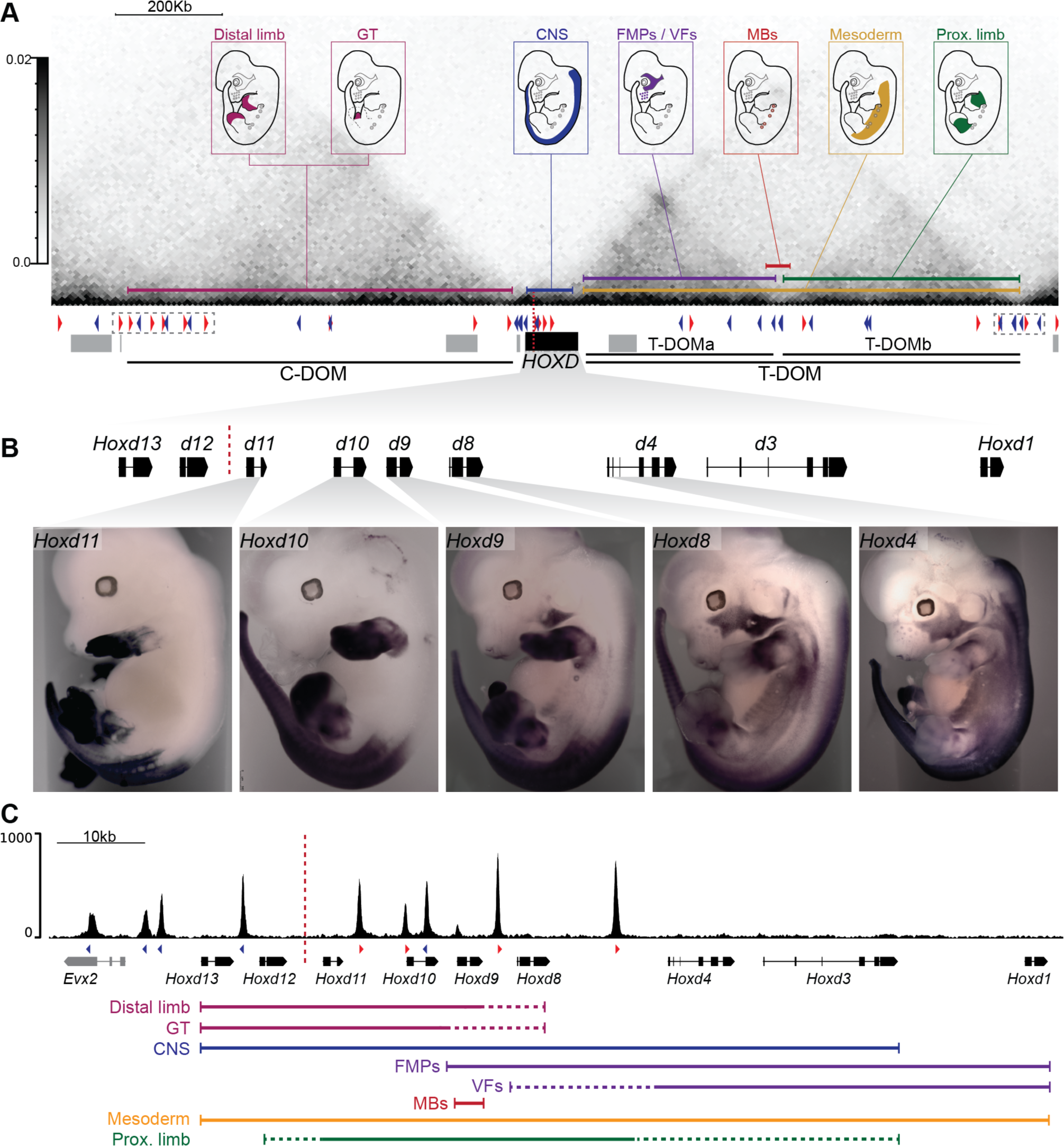
Distribution of CTCF binding sites at the *HoxD* locus and pleiotropic regulation. **A)** Capture Hi-C profile of E9.5 trunks covering the *HoxD* cluster and flanking TADs (T-DOM and C-DOM)(mm10: chr2:73779626-75669724). The embryos on top illustrate various sites of *Hoxd* gene expression and are linked to coloured horizontal bars indicating the positions of the related enhancer sequences, either within the cluster itself (black box) or in the flanking TADs. Dashed grey boxes highlight dense arrays of CBS at TAD borders. GT: genital tubercle; CNS: central nervous system; FMPs: facial muscle progenitors; VFs: vibrissae follicles (whisker pads); MB: mammary buds. The positions and orientations (red or blue) of CTCF binding sites (triangles) are indicated below the Capture Hi-C heatmap. Surrounding genes are shown as filled grey boxes. **B)** WISH analysis showing the expression of *Hoxd11, Hoxd10*, *Hoxd9*, *Hoxd8*, and *Hoxd4* in E12.5 wildtype embryos, to highlight the various overlapping expression patterns for subsets of these genes across distinct embryonic structures. **C)** CTCF ChIP-M profile at *HoxD* using E10.5 trunks (mm10: chr2:74650810-74767377). CBSs are shown as red and blue arrowheads as in (A). The dashed red line separates the two arrays of CBSs showing divergent orientations. The coloured lines on bottom indicate the subsets of *Hoxd* genes expressed in any given embryonic structures. The colours correspond to those delineating the related enhancers in panel (A).

For each of these regulations, a distinct -though overlapping- set of contiguous *Hoxd* genes are selected as targets. In every case, the subgroup of genes responding to a given regulation is delimited by pairs of CTCF sites showing the same orientation, towards the TAD where the related enhancers are localized (Fig. 1). For example, digit enhancers located within C-DOM control *Hoxd13* to *Hoxd10,* whereas forearm enhancers present in T-DOM mostly contact the *Hoxd11* to *Hoxd9* DNA interval. In both cases, the corresponding H3K27ac profiles over these different sets of target genes are delimited by different pairs of occupied CTCF sites (Rodriguez-Carballo et al., 2017).

This dense series of CTCF sites is spread around 50 to 60 Kb, with sites mostly found in between transcription units, a distribution that appeared conserved amongst the four *Hox* mammalian clusters as well as between tetrapod species (Yakushiji-Kaminatsui et al., 2018). This is suggestive of a strong selective pressure to maintain such an organization and thus of potentially important functions for these sites. Indeed, previous studies have revealed the role of individual CBSs in determining micro-boundaries as exemplified with the *HoxA* and *HoxC* clusters (Ghasemi et al., 2021; Luo et al., 2018; Narendra et al., 2015, 2016a; Su et al., 2021). In particular, Narendra et al. showed that the deletion of such sites would modify the extent of *Hox* genes expressed during the formation of the major body axis (in particular in motoneurons cultures) and hence that these sites may behave as micro-insulator elements between neighboring genes (Narendra et al., 2015, 2016a). However, besides the formation of the main body axis, the potential function of CTCF sites in the complex interactions between the large flanking regulatory landscapes and the various subsets of target genes remained to be determined.

In this study, we used a cumulative *in-cis* CRISPR/Cas9 genome editing strategy to disrupt the five CBSs located on one side of the TAD boundary, i.e., within the anterior and central part of the *HoxD* gene cluster. We report the analysis of mouse lines either carrying single mutated sites or the full series of combined mutations. This progressive allelic series starts with the most ‘anteriorly’ located (closer to T-DOM) CBS and follows with the first two, three, four and five CBSs *in cis*, the latter combination removing all those CTCF sites located on the telomeric side of the TAD border. We analyzed the impact of these various mutations, both on *Hoxd* gene expression and chromatin architecture across different tissues, and described associated patterning defects along the major body axis. We conclude that CTCF sites within *Hox* clusters are important for the capacity of remote enhancers to select sub-groups of target genes. However, not all CTCF sites share the same functional task and, while some sites appear to behave as insulators, others seem to have an opposite, anchoring capacity. Notably, some CBSs can display both activities in different tissues. Also, while the selective removal of all CTCF and RAD21 binding on one side of the TAD border certainly resulted in increased inter-TAD interactions, a TAD boundary was still clearly present, indicating that the series of remaining CTCF sites with the opposite orientation was sufficient to maintain the opposite tropism in enhancer-promoter interactions, even though it was weaker and less precise.

## RESULTS

### Evolutionarily conserved CTCF sites in *Hox* clusters

During gastrulation, *Hox* genes are transcribed along the neural tube, paraxial and lateral mesoderm with expression boundaries that reflect their respective positions within the cluster (Gaunt et al., 1988) (Fig. 1). Subsequently, various subsets of these genes are transcribed across a number of embryonic structures, as exemplified by the developing limbs where digit- and forearm enhancers, located in opposite TADs, regulated partially overlapping subgroups of *Hoxd* genes (Fig. 1A) (Andrey et al., 2013). In another context, *Hoxd1* to *Hoxd4* and *Hoxd1* to *Hoxd9* are expressed in the emerging vibrissae follicles (VFs) and in facial muscle progenitors (FMPs), respectively, driven by enhancers located within the T-DOM (Hintermann et al., 2021)(Fig. 1A-C). Alternatively, a single gene can display one particular functionality such as *Hoxd9,* which is the only *Hoxd* gene expressed in the mesenchymal condensates of the future embryonic mammary glands (Chen and Capecchi, 1999), an expression controlled by an enhancer also located within T-DOM, at the boundary between two sub-TADs, T-DOMa and b (Fig. 1A-C) (Schep et al., 2016).

To address whether bound CTCF could be instrumental in the selection of distinct target promoters by such remote enhancers, we looked at CTCF occupancy over the 2 Mb large *HoxD* landscape, together with global chromatin interaction profiles derived from a capture Hi-C approach (Fig. 1A, C). We scored nine occupied CTCF sites in the *HoxD* cluster and over the immediately adjacent *Evx2* gene (Fig. 1C) and two dense arrays of nine and seven CTCF peaks at the opposite borders of the C-DOM and T-DOM, respectively (Fig. 1A, dashed boxes). In contrast, CTCF peaks were less densely distributed within the latter two regulatory landscapes, though with a higher number in T-DOM than in C-DOM. Also, a comparison of CTCF occupancy across several embryonic structures revealed that CTCF binding over the entire *HoxD* genomic landscape was largely comparable in all tissues analyzed (Supplemental Fig. S1), as observed genome-wide (Phillips-Cremins et al., 2013; Schmidt et al., 2013).

Motif analysis revealed that the orientations and positions of the various CBSs tightly correlated with the observed topology of interactions across these domains (Fig. 1A). For instance, within the sub-TAD T-DOMa, two divergent CBSs delimit a domain preferentially interacting with the *Hoxd1* gene, while the sub-TAD boundary region is enriched in CBSs with a negative orientation, defining the extent of the *Hoxd3*-*Hoxd8* preferential interactions (Fig. 1A). Furthermore, the CBSs located either in the centre of C- DOM or within T-DOMa and b correlate well with the extent of local interaction domains within their respective higher-order structures (Fig. 1A).

Within the *HoxD* cluster itself, CBSs are arranged in two arrays of motifs with divergent orientations. All but one CBSs within the anterior and central portion of the cluster (from *Hoxd1* to *Hoxd11*) display a forward orientation and face convergent CBSs located in T-DOM, which are for the most present in the reverse orientation. Instead, CBSs mapping between *Hoxd12* and *Evx2* have a reverse orientation, coinciding with their interaction with CBSs located within or at the centromeric border of C-DOM (Fig. 1C). As expected, the inversion in the orientations of CBSs within the *HoxD* cluster matches the TAD boundary in many cell types, even though this boundary was shown to be able to slightly shift over the *Hoxd11* to *Hoxd10* genes in particular circumstances (Andrey et al., 2013) (Fig. 1). Of note, this organization of the mouse *HoxD* cluster is largely maintained amongst the four *Hox* mammalian clusters (Supplemental Fig. S2), as well as in both the human and chicken orthologous *Hoxd* gene clusters (Supplemental Fig. S3) and is even similar to what was reported in squamates (Guerreiro et al., 2016). Therefore, the presence, distribution and orientation of CBSs within *Hox* clusters is globally conserved across tetrapods, supporting the idea that they may importantly contribute to some aspects of their regulation during development.

In addition, the analysis of RAD21 occupancy, a structural component of the Cohesin complex, (Cheng et al., 2020) and references therein) revealed a substantial enrichment in several of these CBSs, either within the *HoxD* cluster or in the flanking TADs. Amongst the formers, RAD21 accumulation was maximal at CBS1, 2 and 4, as well as in CBS6 to 8. Instead, weak or no RAD21 enrichment was observed in the centrally-located CBS3 and 5 or at CBS9 (Fig. 2A). While this pattern of accumulation of RAD21 is compatible with the corresponding CTCF sites being involved in long-range interactions through loop formation, the heterogeneity in the RAD21 profile, at least in this tissue, suggests that not all occupied CTCF sites may share the exact same function.

**Figure 2:**
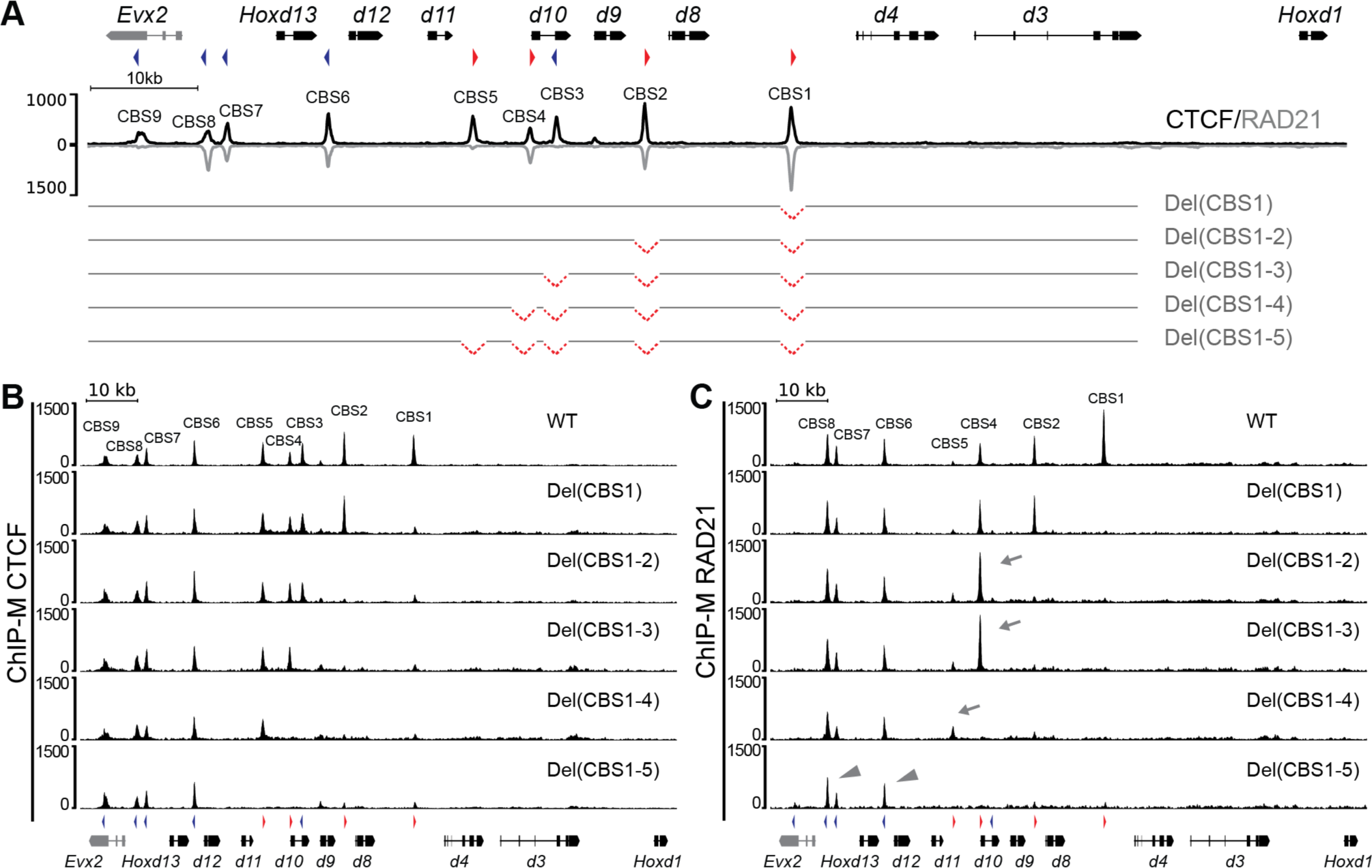
The *in-cis* CTCF mutant allelic series. **(A)** CTCF (black) and RAD21 (grey) ChIP-M profiles at control and mutant *HoxD* loci, using dissected E10.5 trunks (mm10: chr2:74650810-74767377). The CTCF binding site (CBS) number is indicated above the peaks with its orientation on top. A schematic representation of the CBS deletion alleles is shown below, only indicating the combined mutants *in-cis*. Single CBSs mutations are not shown except for Del(CBS1). (**B-C)** CTCF (**B**) and RAD21 (**C**) ChIP-M enrichments at the *HoxD* locus in E10.5 trunks of either control (WT), or Del(CBS1), Del(CBS1-2), Del(CBS1-3), Del(CBS1-4) or Del(CBS1-5) homozygous embryos. No significant difference in enrichment over the non-mutated CBSs was scored and no cryptic binding sites was revealed. For RAD21, grey arrows indicate distinctive changes in RAD21 accumulation in some mutant alleles, whereas arrowheads highlight the stable enrichment of RAD21 over C-DOM oriented CBSs (blue triangles), independently of any mutation (mm10: chr2:74650810-74767377).

### An *in cis* allelic series for mice mutant for CTCF binding

To investigate the potential function of these CBSs either in the organization of the TAD boundary or during the function of remote enhancers, we generated an allelic series of mutant mice carrying homozygote micro-deletions of all those CBSs located at the TAD border, on the T-DOM side (Fig. 2A, CBS1 to CBS5; Supplemental Fig. S4). We designed sgRNAs targeting the various CBSs identified within the ChIPmentation (ChIP-M) CTCF peaks (Supplemental Fig. S4, see Material and Methods) and co-electroporated them in fertilized mouse oocytes together with the Cas9 mRNA. After control by Sanger sequencing, F0 animals were crossed to produce stable mutant lines (Fig. 2A,Supplemental Fig. S4). In this way, we generated from 6bp to 78bp long micro-deletions impacting the predicted CBSs (Supplemental Fig. S4B) and affecting those nucleotides required for CTCF binding (Hashimoto et al., 2017; Lobanenkov et al., 1990).

To produce the series of micro-deletions *in-cis*, we first obtained separately animals homozygous for the individual deletion of the CBS1, referred to as *HoxD^Del(CBS1)-/-^* or Del(CBS1) and the CBS2 (*HoxD^Del(CBS2)-/-^*) or Del(CBS2). We then electroporated the sgRNA targeting CBS1 into zygotes heterozygous for Del(CBS2) and recovered the double mutant *in cis HoxD^CBS(1-2)-/-^* or Del(CBS1-2). This operation was reiterated three times by using the various sgRNAs on the newly produced strains carrying mutations *in cis* to eventually obtain the *HoxD^CBS(1-5)-/-^* mice or Del(CBS1-5), where the five contiguous CTCF sites located on the T-DOM side of the TAD boundary were mutated on the same chromosome. For each mutant, and before processing to the next mutation *in cis*, we assessed CTCF binding by ChIP-M in the post occipital region of E10.5 wildtype and homozygous mutant embryos. As expected, these mutations mostly abolished CTCF binding to the target sites (Fig. 2B). Of note, and in contrast to what was reported for CBS mutagenesis in other genomic contexts (Narendra et al., 2016a; Paliou et al., 2019), we did not observe any cryptic CTCF binding site, which would have been revealed after the mutation of neighbouring sites. Also, the binding enrichments observed at the remaining CTCF sites were not modified (Fig. 2B).

The effect of these micro-deletions on CTCF binding was also verified by performing ChIP-M for RAD21 using wildtype and various mutant embryos (Fig. 2C). AS expected, the disruption of the CTCF motifs resulted in the loss of RAD21 at the targeted CBSs. Furthermore, in the mutant conditions, RAD21 was redistributed and increased accumulations were observed at CBSs located next to the deleted site(s). For example, RAD21 accumulation was increased at CBS4 in the Del(CBS1-2) mutant, an increase reinforced in the Del(CBS1-3) allele (Fig. 2C, arrows; Supplemental Fig. S5). Changes in RAD21 enrichment were also observed at CBS5, a CTCF site that accumulates virtually no RAD21 in the wildtype condition (Fig. 2C, arrow; Supplemental Fig. S5). Yet no difference in RAD21 enrichment was observed neither at CBS6 to 9, nor at CBS3, all these CTCF sites displaying a reverse motif orientation (Fig. 2C, arrowheads; Supplemental Fig. S5), further supporting an involvement of CBS1, 2, 4 and 5 in forming chromatin structures with CTCF sites located further telomeric, within T-DOM.

### Impact of CTCF binding sites deletions *in cis* upon chromatin topology

To assess the impact of these progressive deletions of CBSs upon the global TAD architecture, we performed Capture Hi-C over the region covering the *HoxD* cluster and flanking TADs. We used trunks of E9.5 control embryos, where *Hox* genes are transcribed, and compared the interaction profiles with those obtained from Del(CBS1), Del(CBS1-3) and Del(CBS1-5) homozygous fetuses. The *HoxD* cluster is positioned between the two C-DOM and T-DOM TADs, which are insulated from one another by the CTCF-dependent boundary present within the gene cluster (Fig. 3A) (Rodriguez-Carballo et al., 2017). In this embryonic material, the TAD-separation score using the hicFindTADs algorithm identified the position of this boundary between *Hoxd13* and *Hoxd12* (Fig. 3A, Supplemental Fig. S6C).

**Figure 3:**
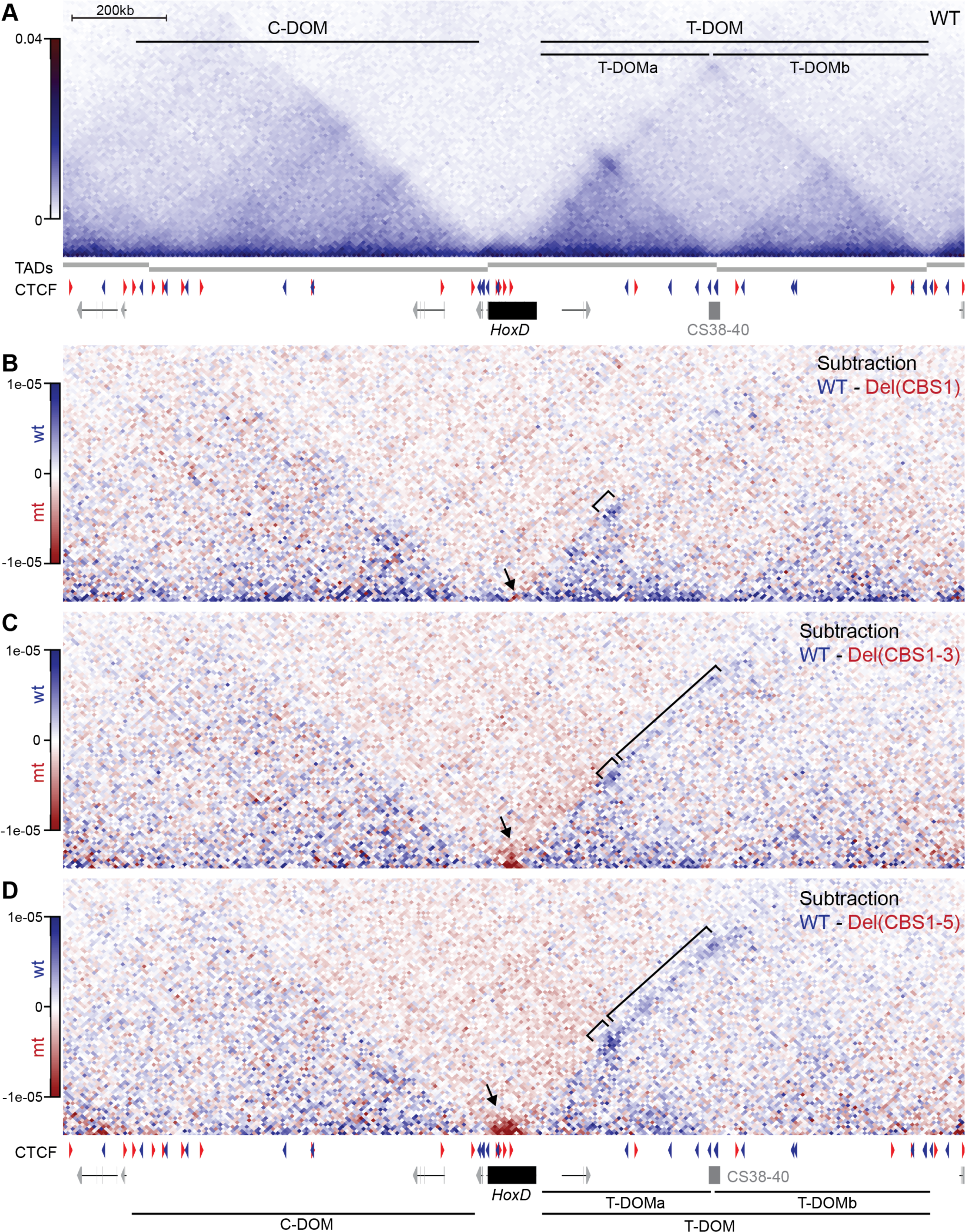
Changes in chromatin topology in CBS mutants *in vivo*. **(A)** Capture Hi-C map of control (WT) E9.5 trunks with the *HoxD* cluster (black rectangle) and flanking TADs (C-DOM and T-DOM) (mm10: chr2:73779626-75669724). TAD or sub-TADs (T-DOMa and T-DOMb) were identified using the hicFindTADs algorithm (window size of 240kb) and are represented by grey bars below the heatmap. Red and blue arrowheads below indicate the orientation of the CBSs. The sub-TAD boundary region CS38-40 is shown as a grey box. **(B-D)** The subtraction maps between the cHi-C profiles from control versus Del(CBS1), Del(CBS1-3) and Del(CBS1-5) homozygous embryos are displayed, with blue bins pointing to chromatin interactions that are more prevalent in control cells, while red bins represent interactions enriched in mutant alleles The black arrows indicate the increase of intra-cluster interactions in mutant alleles. The small brackets point to a progressive decrease in interactions between the gene cluster and the CS38-40 region in the mutant alleles, whereas the large brackets indicate the loss of contact frequency between the *HoxD* cluster and T-DOMb in both the Del(CBS1-3) and Del(CBS1-5) alleles. C-DOM, T-DOM, T-DOMa, and T-DOMb are represented below as in panel A (mm10: chr2:73779626-75669724).

For each mutant condition, a subtraction was carried out from the control interaction profile and the various subtractions with gained interactions in red and lost interactions in blue were compared (Fig. 3B-D). The abrogation of CBS1 resulted in two slight yet significant changes: First, an increase of self-interactions was observed between the *Hoxd1 to Hoxd8* genes (Fig. 3B, arrow; Supplemental Fig. S6A, B), without altering the position of the TAD border (Supplemental Fig. S6C). Secondly, a decrease in interactions between the gene cluster and the CS38-40 region (Fig. 3B, bracket), a region containing three CTCF sites oriented towards the *HoxD* cluster, which acts as a sub-TAD boundary within the T-DOM (Fig. 3A). An average signal quantification and virtual Capture-C profiles using *Hoxd4* as a viewpoint confirmed this reduction (Supplemental Fig. S7A-C).

These two differences observed in the interaction profile of Del(CBS1) were strongly reinforced when the capture Hi-C profile of the Del(CBS1-3) was subtracted from the control counterpart. Indeed, a marked increase in intra-cluster interactions was scored, which extended up to *Hoxd10* (Fig. 3C, arrow; Supplemental Fig. S6A, B). Furthermore, the TAD-separation score analysis revealed a change in border position that was now called at a more centromeric position, after the *Evx2* gene (Fig. 3C; Supplemental Fig. S6C). Secondly, the frequencies of interactions between the gene cluster and T-DOM not only showed a loss in contacts with the CS38-40 region (Fig. 3C, small bracket), but also throughout the most telomeric sub-TAD T-DOMb (Fig. 3C, large bracket; Supplemental Fig. S7A, B). To facilitate data visualization, we generated virtual Capture-C profiles using different viewpoints. This analysis revealed a loss of interactions between the *Hoxd4* promoter and the CS38-40 region, as well as a reduction in contacts between the cluster and the T-DOMb, when using either *Hoxd9* or the T-DOMb 3’border as viewpoints (Supplemental Fig. S7C).

This effect was again enhanced in the Del(CBS1-5), i.e. after the deletion of all CTCF sites oriented towards the T-DOM (Fig. 3D). The intra-cluster interactions were strengthened and extended up to *Hoxd13,* (Fig. 3D, arrow; Supplemental Fig. S6A, B), with a TAD boundary being called after the *Evx2* gene (Fig. 3D; Supplemental Fig. S6C). In addition, the interactions with the CS38-40 region were further lost, when compared to the Del(CBS1-3) condition. Similarly, the decrease in global contact frequency between the *HoxD* cluster and the sub-TAD T-DOMb was further enhanced (Fig. 3D, large bracket; Supplemental Fig. S7A, B), with a loss of interactions even observed with the T-DOM telomeric TAD border. These results were confirmed by virtual Capture-C, when using various viewpoints (Supplemental Fig. S7C). Therefore, the progressive deletions of intra-cluster CTCF sites clearly released long-range interactions between the gene cluster itself and various parts of T-DOM, where the cluster is normally anchored. Likely as a consequence of this lack of long-range contacts, the cluster became more compact and thus increased its local interactions. Finally, none of the mutant interaction profiles showed any major differences in the contacts between the gene cluster and the C-DOM.

### Impact of alterations in chromatin structure upon *Hoxd* genes expression

T-DOM contains several enhancers necessary for the proper expression of *Hoxd* genes and hence we assessed whether these alterations in contact distribution observed in the various CBS deletion alleles were paralleled by either quantitative or qualitative modifications in gene expression. We performed whole-mount RNA *in situ* hybridization (WISH) across our mutant lines and their control littermates and examined gene expression along the major body axis. While expression in the spinal cord appears to be regulated by intra-cluster control sequences (e.g. (Tschopp et al., 2012) transcription in mesoderm derivatives is at least partly regulated by elements located within T-DOM (Fig. 4A, brown). In Del(CBS1) mutant fetuses, we observed an anteriorization of the expression domain of *Hoxd8,* the gene positioned just 5’ of the deleted CTCF binding site (CBS1)(Fig. 4B), whereas the expression domains of both *Hoxd4* and *Hoxd9* remained unaltered (Supplemental Fig. S8A, B). In the Del(CBS1-2) allele, we observed an anteriorization of the domain of expression of *Hoxd9* (Fig. 4B) without affecting the expression of *Hoxd10* (Supplemental Fig. S8C). In turn, the domain of expression of the latter gene was anteriorized in Del(CBS1-3) mutant fetuses (Fig. 4B), while *Hoxd11* transcripts remained indistinguishable from control littermates (Supplemental Fig. S8D). In both Del(CBS1-4) and Del(CBS1-5) homozygous embryos, the *Hoxd10* and *Hoxd11* expression domains were anteriorized too (Fig. 4B; Supplemental Fig. S8F). Although to a lesser extent, the expression domain of more anterior genes was still affected in these alleles, as exemplified by *Hoxd9* staining in the Del(CBS1-4) (Supplemental Fig. S8E).

**Figure 4:**
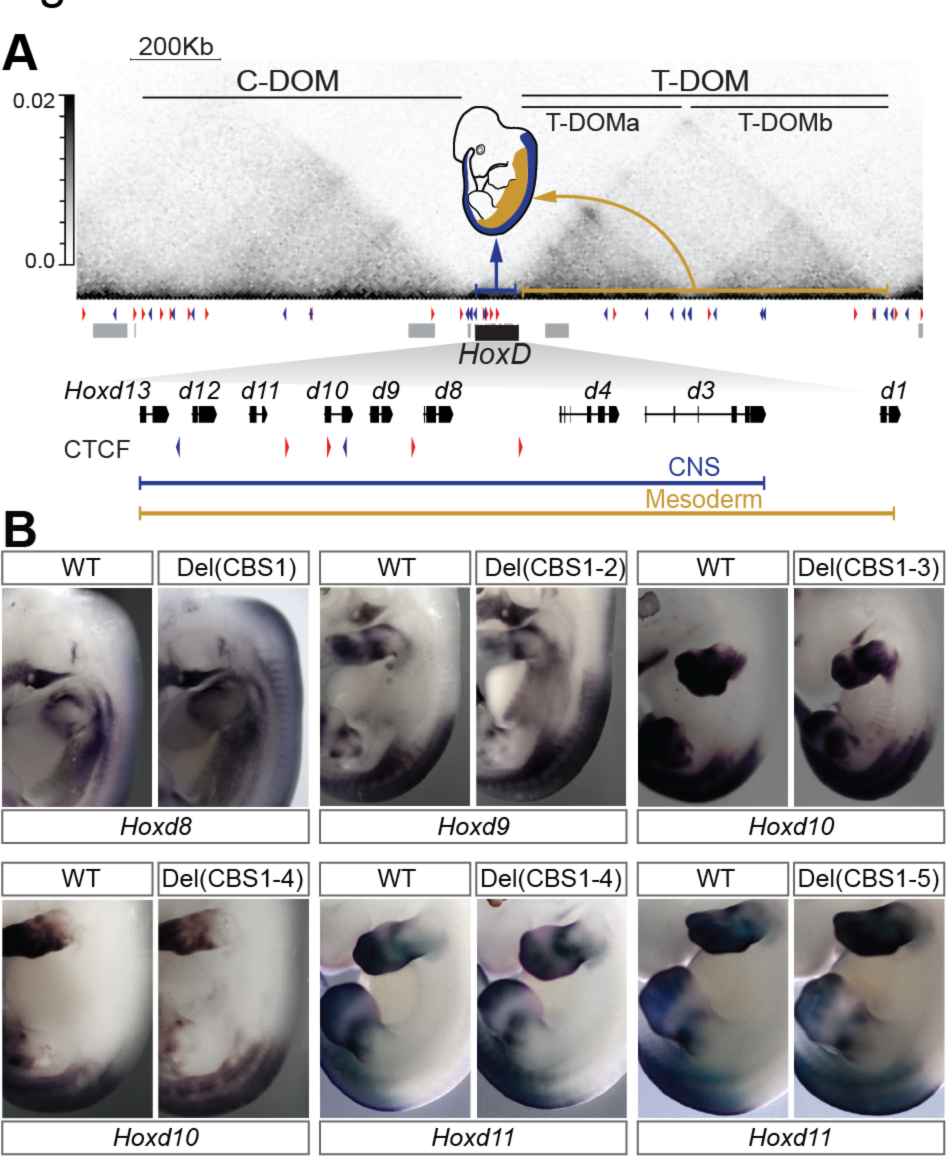
Alterations in *Hoxd* genes expression along the anterior to posterior axis. **A)** CHi-C map from control E9.5 dissected trunk region showing the *HoxD* cluster and neighboring regulatory domains T-DOM and C-DOM (mm10: chr2:73779626-75669724). The schematized embryo on top of the heatmap represents the regulation of *Hoxd* gene expression in the spinal cord (CNS; blue) by enhancer elements located within the gene cluster (blue arrow), whereas gene transcription in axial mesoderm (yellow) is in large part controlled by regulatory elements located in the T-DOM (yellow arrow). The positions of the *HoxD* cluster (black) and the surrounding genes (grey) are shown below. On the bottom, a zoom-in of the *Hoxd* genes and CBS is shown. The blue and yellow lines underline which *Hoxd* genes are expressed either in the CNS, or in the mesoderm, respectively. **(B)** WISH analysis showing the expression of different *Hoxd* genes in the Del(CBS1), Del(CBS1-2), Del(CBS1-3), Del(CBS1-4) and Del(CBS1-5) E12.5 embryos with corresponding control littermates. An anteriorization of expression in the trunk was generally observed for the gene positioned immediately 5’ to the deleted CBS.

Changes in axial patterning due to alterations in the expression of *Hox* genes have been widely described (see (Carapuço et al., 2005; Kessel and Gruss, 1991a; Mallo et al., 2010). We performed micro-CT scans of control and homozygous mutant adult mice skeletons and, as expected, we observed alterations in the spine of mutant animals, associated with changes in expression pattern (Supplemental Table S1). For example, 36 percent of Del(CBS1-4) mutant mice have four lumbar vertebrae (L4), instead of the L5 condition observed in this genetic background (Supplemental Table S1). Such lumbo-sacral defects were previously associated with alterations in *Hoxd11* expression (Gerard et al., 1996; Wellik and Capecchi, 2003; Zakany et al., 1996). Noteworthy, this morphological alteration was both less severe and less prevalent in Del(CBS1-5) animals (Supplemental Table S1).

More than half of the mutants carrying any of the three genotypes analyzed displayed defects at the atlanto-occipital junction, involving the basioccipital bone, the atlas, and the axis (Supplemental Table S1). A minor proportion of both Del(CBS1-3) and Del(CBS1-4) mutant animals also showed the asymmetrical presence of cervical ribs, a malformation at the cervico-thoracic transition (Supplemental Table S1). These results confirm the physiological relevance of CTCF-mediated regulation of *Hoxd* gene expression along the A-P body axis and are in agreement with previous work involving the *HoxA* and *HoxC* clusters, where the local removal of CTCF led to such transformations (Narendra et al., 2016a).

### Differential impacts of CBS mutations upon *Hoxd* gene regulation in mesoderm derivatives

The expression of *Hoxd1* to *Hoxd4* in the dermal papilla of vibrissae follicles (VFs) depends on CREs located within a region of the sub-TAD T-DOMa that preferentially interacts with *Hoxd1* and the most ‘anterior’ part of the *HoxD* cluster (Fig. 5A). Likewise, a set of enhancers located across the same T- DOMa drives the expression of *Hoxd1* to *Hoxd9* into facial migrating muscle progenitors (FMPs) (Hintermann et al., 2021)(Fig. 5A). We asked whether the disruption of intra-cluster CBSs would affect the various distributions of target genes for these two regulatory specificities, which concern either genes located within the part of the gene cluster that is devoid of CTCF sites, or genes extending slightly behind CBS1 and 2. In Del(CBS1) mutant embryos, *Hoxd8* was upregulated in both VFs and FMPs, when compared to control littermates (Fig. 5B, arrowhead and asterisk) in contrast to *Hoxd9,* which was not affected (Supplemental Fig. S9A). Instead, *Hoxd9* was ectopically activated in these structures in Del(CBS1-2) mutant fetuses (Fig. 5B, arrowhead and asterisk), while *Hoxd10* remained inactive in these mutants (Supplemental Fig. S9A). Noteworthy, a tissue-specific effect of the CBS deletions upon *Hoxd* genes expression was observed. Indeed, while the deletion of CBS1 and 2 allowed for the extension of the gene subset expressed in VFs up to *Hoxd9*, the additional mutation of CBS3 triggered the transcription of *Hoxd10* in the FMPs, yet not in the VFs (Fig. 5B, asterisk). No changes in the transcription domains of *Hoxd* genes located 3’ to the mutated CBSs was observed, as exemplified by the analysis of *Hoxd1* and *Hoxd4* transcript distribution in the Del(CBS1) and Del(CBS1-2) mutants, respectively (Supplemental Fig. S9B). Also, we were unable to observe any changes in the expression of *Hoxd9 to Hoxd11* in the Del(CBS1-5) mutant embryos (Supplemental Fig. S9C).

**Figure 5:**
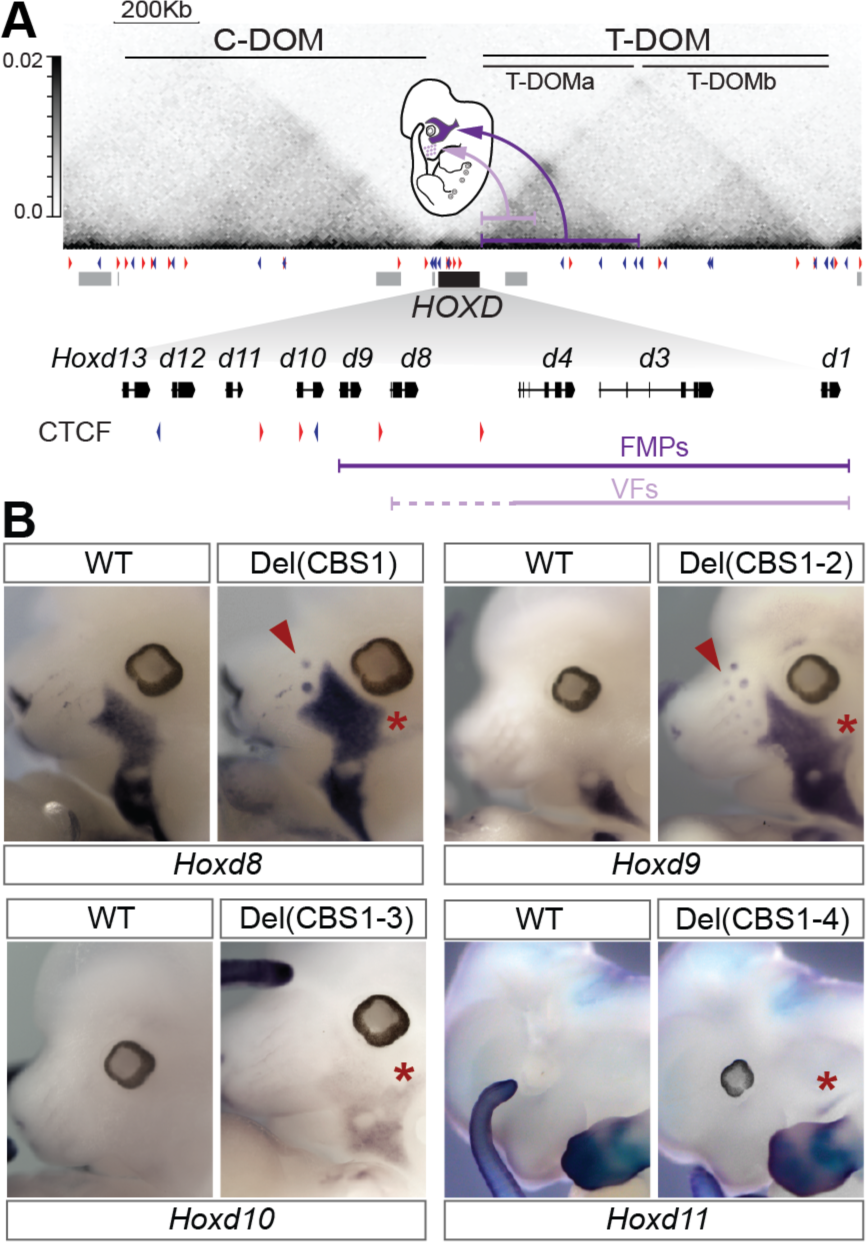
Alterations in *Hoxd* genes expression in facial structures. **(A)** Schematic representation of *Hoxd* gene regulation in both the vibrissae follicles (VFs, light purple) and some facial muscle precursors (FMPs, dark purple) as well as the DNA segments within T-DOM where the corresponding enhancers are located (Hintermann et al., 2021). A magnification of the gene cluster is shown below with the subsets of *Hoxd* genes expressed in each structure. CBS and their orientations are indicated as red or blue triangles (**B)** WISH showing *Hoxd* gene expression in the VFs and FMPs of control (WT) or various CBS deletion alleles, as indicated in each panel. Red arrowheads and asterisks indicate either an upregulation or an ectopic expression in the VFs and FMPs of each mutant allele, respectively.

In summary, the disruption of the intra-cluster CBSs generally led to the ectopic transcriptional activation of the gene located immediately centromeric to the deleted CBSs, suggesting that, in these contexts, CTCF sites operate as insulator elements constraining gene-enhancer interactions such as to delimit specific subsets of contiguous genes competent to respond to (a) particular remote enhancer(s). The upregulation of *Hoxd9* and *Hoxd10* in the FMPs was considerably weaker, if not absent, from Del(CBS1-4) and Del(CBS1-5) mutant embryos, in contrast to the Del(CBS1-2) and Del(CBS1-3) conditions (Fig. 5B; Supplemental Fig. S9C).). This, together with the loss of *Hoxd11* ectopic activation in Del(CBS1-5) embryos, suggests that CBS4 and CBS5 may have an anchoring function for the FMPs and VF enhancers and that their mutations abrogate the gains of expression observed upon deletions of CBS1 to CBS3, the latter sites being used as insulators in these contexts.

Finally, mesoderm cells of the embryonic mammary bud contain high levels of *Hoxd9* mRNAs, in contrast to all other *Hoxd* genes, which are downregulated in this structure around E12 to E13 (Schep et al., 2016). In these cells, *Hoxd9* is controlled by a eutherian-conserved enhancer located within the CS38-40 sub-TAD boundary within the T-DOM (Supplemental Fig. S9D). We looked at the effect of disrupting CBS2 in isolation, i.e., the CTCF site located between *Hoxd8* and *Hoxd9* (Supplemental Fig. S9E) and the analysis of Del(CBS2) mutant embryos revealed an upregulation of *Hoxd9* in paraxial and lateral mesoderm, yet considerably weaker than that observed in Del(CBS1-2) mutant mice (Fig. 4B; Supplemental Fig. S9F). However*, Hoxd9* transcription in the mammary bud mesenchyme was no longer detected (Supplemental Fig. S9F) and *Hoxd10* was not ectopically activated in the mammary bud mesenchyme nor in the main body axis of Del(CBS2) embryos (Supplemental Fig. S9G). These results indicate that CBS2 is required for the proper anchoring of the mammary bud enhancer with its target gene, while it operates as an insulator element in the main body axis. Together with the alterations observed in the VFs and FMPs, these data indicate that the disruption of CBS differentially impacts *Hoxd* gene transcription across tissues, with effects that cannot be solely attributed to their role as insulator elements.

### Effects of CBS disruptions upon *Hoxd* gene regulation in developing proximal limb buds

During limb buds development, *Hoxd* genes are regulated in a bimodal manner by the two TADs, with *Hoxd9* to *Hoxd11* initially under the control of forearm T-DOM enhancers, whereas *Hoxd10* to *Hoxd13* become subsequently controlled by C-DOM enhancers in digit cells (Figs 1A, 6A and 7A) (Andrey et al., 2013; Beccari et al., 2016; Rodriguez-Carballo et al., 2017). To evaluate the function of CBSs in the definition of these subsets of target genes, as well as upon the 3D chromatin conformation in limb bud cells, we performed RNA-sequencing (RNA-seq) and Capture Hi-C on micro-dissected proximal and distal E12.5 forelimb cells derived from both our mutant CBS allelic series and control embryos.

**Figure 6:**
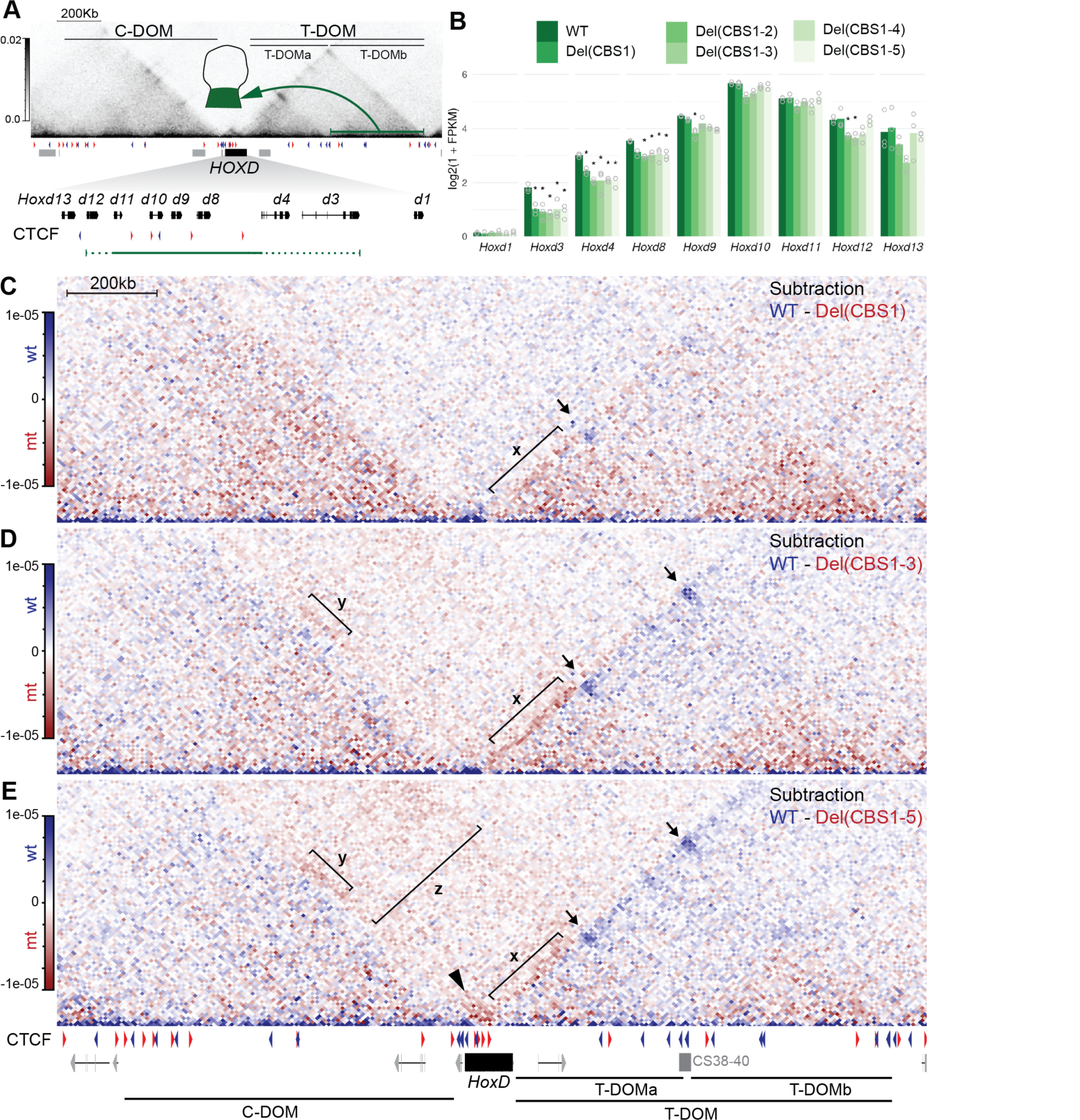
Impact of the CBS alleles upon *Hoxd* gene regulation in developing proximal limb buds. **(A)** Capture Hi-C map of control E12.5 dissected proximal forelimb (PFL) cells with the *HoxD* cluster (black rectangle) and the neighbouring T-DOM and C-DOM (mm10: chr2:73779626-75669724). The schematics limb shows the T-DOM regulation in proximal cells (green) as a result of multiple enhancers located within T-DOMb (green line and arrow). Below is a magnification showing the CBSs and their orientations (blue and red arrowheads) as well as those *Hoxd* genes responding to this regulation (green line). (**B)** Bar plot representing *Hoxd* genes expression (log2(1+FPKM)) in PFL cells. Hollow circles show the replicates for each sample and the asterisks point to those samples where significant differences in transcript levels were scored between control and mutant alleles (absolute log2 fold change above 0.58, adjusted p-value < 0.05) (**C-E)** Capture Hi-C subtraction maps of control (blue) and mutant (red) E12.5 PFL cells. The subtracted alleles are indicated on the upper right corners. The arrows represent the incremental loss of interactions between both the sub-TAD border region CS38-40 and the telomeric T- DOMb border region, and the part of the *HoxD* cluster where various CBSs were deleted in the mutant alleles. Bracket x indicates an increase in interactions between *HoxD* and T-DOMa, stronger in the Del(CBS1-3) than in the Del(CBS1-5). In contrast, intra-cluster interactions are stronger in the Del(CBS1-5) than in Del(CBS1-3)(arrowhead in (**E**). Bracket y highlights a gain in interactions between the ‘anterior’ genes and the centromeric end of C-DOM in Del(CBS1-3) and Del(CBS1-5). Bracket z points to a general increase in interactions between T-DOMa and a region rich in CTCF sites at the C-DOM centromeric border in Del(CBS1-5).

Using the RNA-seq datasets, we assessed the effects of various CBS deletions first by performing a principal component analysis (PCA) considering the expression levels of the 500 most variant autosomal protein-coding genes. As expected, principal component 1 (PC1), which explained 83% of the total gene expression variance, separated the distal forelimb from the proximal forelimb (Supplemental Fig. S10A). Along PC2, which accounted only for 5% of the total variance, we observed that samples from the same litter tend to cluster together, illustrating a “litter effect” on the set of samples (Supplemental Fig. S10A). These observations were corroborated by the expression clustering based on pairwise Euclidean distances between samples (Supplemental Fig. S10B). These data indicated that the various CBS deletions have a negligible impact on the overall transcription profiles of the developing limbs, in agreement with the lack of major alterations in limb morphology in mutant animals throughout the allelic series.

We then conducted pairwise differential gene expression analyses (absolute log2 fold change above 0.58, adjusted p-value < 0.05) of the proximal forelimb samples of control and various CBS mutant alleles. Using these parameters, we observed between 6 and 195 protein-coding genes differentially expressed in PFL (Supplemental Fig. S11A; Supplemental File 1), thus confirming the weak differences across samples observed in the PCA. However, the expression levels of anterior *Hoxd* gene (*Hoxd3, Hoxd4, Hoxd8*) in proximal forelimb (PFL) were consistently decreased in all mutant alleles, with the exception of *Hoxd8* in the Del(CBS1) allele (Fig. 6B). These differences could not be attributed to the litter effect and the intersection of differentially expressed genes identified *Hoxd3* and *Hoxd4* as being the only mis-regulated genes in all alleles. Similarly, *Hoxd8* was the only differentially expressed gene in the Del(CBS1-2), Del(CBS1-3), Del(CBS1-4), and Del(CBS1-5) alleles (Supplemental Fig. S11A: Supplemental File 1). Finally, WISH analysis of *Hoxd8* transcripts in the Del(CBS1-2), Del(CBS1-3), and Del(CBS1-5) alleles confirmed this decrease in mRNAs levels (Supplemental Fig. S12A).

We next asked whether these changes in expression were due to reallocations in interactions between the genes and their regulatory landscapes. We performed Capture Hi-C in micro-dissected proximal forelimb cells of the Del(CBS1), Del(CBS1-3) and Del(CBS1-5) alleles and compared them with control samples (Bolt et al., 2021). In the Del(CBS1) allele, we observed a minor loss of interaction with the CS38-40 T-DOM sub-TAD boundary and a gain of ectopic interactions with T-DOMa, as seen on the subtraction maps (Fig. 6C). When using *Hoxd4* as a viewpoint in virtual Capture-C profiles, we observed a loss of interactions with the CS38-40 region in Del(CBS1) PFLs, which was not observed when using *Hoxd8* (Supplemental Fig. S13), in agreement with the changes in transcripts levels observed for these two genes. Instead, *Hoxd8* increased its interactions with T-DOMa in the Del(CBS1) allele (Fig. 6C, bracket x; Supplemental Fig. S13A, B), a gain that was reinforced in the Del(CBS1-3) allele (Fig. 6D, bracket x; Supplemental Fig. S13A, B). In the latter allele, a very significant decrease in interactions with the CS38- 40 and the T-DOMb regions was scored (Fig. 6D, Supplemental Fig. S13A, B), correlating with the diminution of *Hoxd8* mRNA levels in the PFL of these mutants.

While virtual Capture-C profiles further confirmed the alteration in the interaction between *Hoxd8* and T-DOM (Fig. 6D, Supplemental Fig. S13C), the Del(CBS1-5) allele behave somewhat unexpectedly since the increase in anterior *Hoxd* interaction with T-DOMa was less pronounced, whereas the loss of interactions with CS38-40, T-DOMb and the TAD boundary region was further accentuated when compared to the Del(CBS1) and Del(CBS1-3) alleles (Fig. 6E; Supplemental Fig. S13A-C). Furthermore, in the Del(CBS1-5) allele, we observed an increase in interactions between the anterior genes (e.g. *Hoxd4, Hoxd8)* and posterior *Hoxd* genes (*Hoxd13* to *Hoxd11*) (Fig. 6E, arrowhead; Supplemental Fig. S13C)

In parallel with their loss of contacts with T-DOMb, anterior genes gained some interactions with the centromeric end of C-DOM (Fig. 6E, bracket y), as if this part of the cluster was now free to follow those contacts normally established between the 5’ C-DOM border and the CTCF sites with opposite orientations remaining in the ‘posterior’ part of the *HoxD* cluster. In fact, the entire T-DOMa seemed to be dragged along by these CTCF sites as suggested by the general increase in interaction between the entire gene cluster and the C-DOM 5’ border (Fig. 6E, bracket z).

### Effects of CBS disruptions upon *Hoxd* gene regulation in developing distal limb buds

The general changes observed in gene expression and chromatin organization in distal forelimb (DFL) samples of our various mutant alleles were somewhat comparable, yet slightly different from those observed in PFL. Differential gene expression analyses identified between 6 and 282 protein-coding genes differentially expressed in distal forelimb cells (Supplemental Fig. S11B: Supplemental File 1). In the Del(CBS1-2), Del(CBS1-3) and Del(CBS1-4) alleles, we scored a mild yet significant decrease in mRNA levels of *Hoxd9, Hoxd10* and *Hoxd11* (Fig. 7B). In agreement with RNA-seq datasets, WISH analysis showed decreased signal intensity of *Hoxd9* and *Hoxd11* in the DFLs of these alleles (Supplemental Fig. S12B). We also observed a recovery in transcript levels in the Del(CBS1-5) allele, both by RNAseq and WISH (Fig. 7B; Supplemental Fig. S12C). Intersections between the Del(CBS1-2), Del(CBS1-3) and Del(CBS1-4) samples identified *Hoxd9, Hoxd10,* and *Hoxd11* as the only commonly differentially expressed genes in these samples (Supplemental Fig. S11B), again highlighting the local effect of these deletions on gene expression.

**Figure 7:**
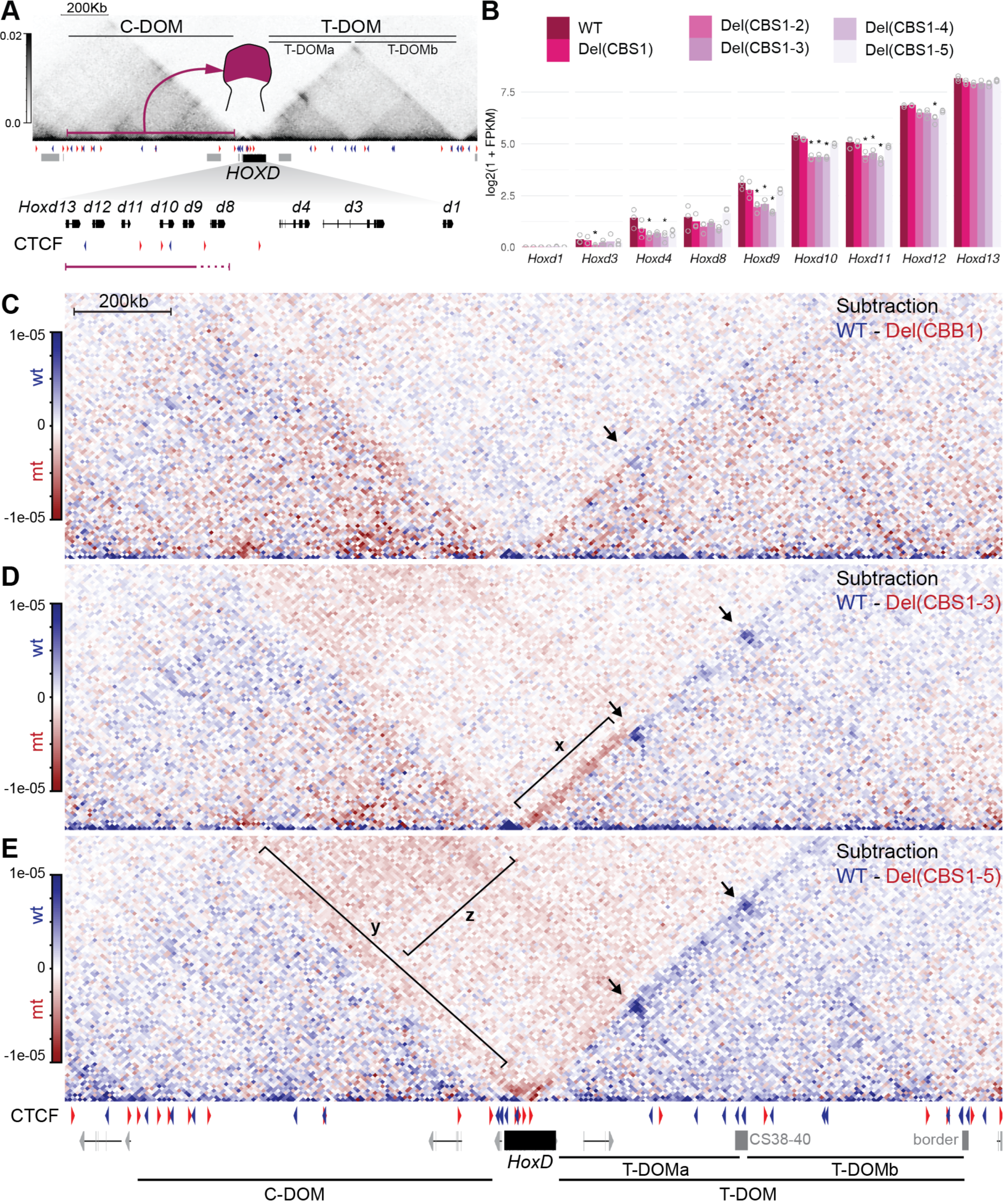
Impact of the CBS alleles upon *Hoxd* gene regulation in developing distal limb buds. **(A)** Capture Hi-C map of control E12.5 dissected distal forelimb (DFL) cells with the *HoxD* cluster (black rectangle) and the neighbouring T-DOM and C-DOM (mm10: chr2:73779626-75669724). The schematics limb shows the C-DOM regulation in distal cells (pink) as a result of multiple enhancers spread over C- DOM (pink line and arrow). Below is a magnification showing the CBSs and their orientations (blue and red arrowheads) as well as those *Hoxd* genes responding to this regulation (pink line). (**B)** *Hoxd* genes expression (log2(1+FPKM)) in DFL. Hollow circles show the replicates for each sample and the asterisks point to those samples where significant differences in transcript levels were scored between control and mutant alleles (absolute log2 fold change above 0.58; adjusted p-value < 0.05). **(C-E)** Subtractions of cHi- C maps between control E12.5 DFL (blue) and various homozygous mutants (red). The subtracted alleles are indicated on the upper right corners. Arrows indicate the loss of interactions, in the mutant alleles, between the ‘anterior’ part of the *HoxD* cluster and both the sub-TAD boundary region CS38-40 and the telomeric T-DOMb border region. Bracket x points to increased interactions between the *HoxD* cluster and T-DOMa in the Del(CBS1-3) mutant **(D)**, less pronounced in the Del(CBS1-5) allele **(E)**. Bracket y indicates increased interactions observed in the Del(CBS1-5) mutant between anterior *Hoxd* genes and C- DOM, whereas bracket z highlights the loss of insulation between C-DOM and T-DOMa, most pronounced in this allele (compare with the Del(CBS1) in **(C)**.

The subtractions of capture Hi-C datasets between DFL and control samples using the Del(CBS1- 2), Del(CBS1-3) and Del(CBS1-5) mutant alleles revealed a progressive loss of interactions between the cluster and the CS38-40 region, T-DOMb and the TAD border regions, similar to what was observed with PFL cells (Fig. 7C-E). Furthermore, we observed a gain of interactions with T-DOMa that was more pronounced in the Del(CBS1-3) allele, which seemed to slightly recover in the Del(CBS1-5) allele (Fig. 7D, E, bracket x). This loss of ectopic interaction with T-DOMa in the Del(CBS1-5) allele may influence the recovery of *Hoxd* gene expression in the Del(CBS1-5) allele.

While the interactions between the C-DOM and the 5’ part of the *HoxD* cluster did not seem to be much affected in the various mutant alleles, contacts were increased between the *Hoxd8* to *Hoxd4* region and C-DOM in the Del(CBS1-5) (Fig. 7E, bracket y), as well as for the entire T-DOMa sub-TAD, as was observed in proximal cells (Fig. 7E, bracket z). However, this gain of contacts was not paralleled by a gain of expression of anterior *Hoxd* genes in digits as revealed by RNAseq and WISH analysis (Fig. 7B and Supplemental Fig. S12A). Overall, these results demonstrate that CBS deletions at the *Hoxd* gene locus result in changes in local chromatin architecture that lead to the downregulation of *Hoxd* gene transcription in proximal and distal forelimbs. However, the observed changes in gene expression indicate a requirement of the intra-cluster CBS to reach a proper level of transcription, rather than for an insulator or anchoring effect related to those observed during the development of the main embryonic body axis.

## DISCUSSION

The disruption or deletion of CTCF-binding sites to study the role of this protein in the organisation of chromatin in 3D and in transcriptional insulation has been reported by using different genetic loci and experimental models. For example, the deletion of some CTCF sites at the mouse *Shh* locus weakened the formation of large chromatin loops between the *Shh* gene and its ZRS enhancer (Paliou et al., 2019). Also, the importance of multiple CTCF sites for proper insulation at TAD borders through the formation of local chromatin domains has been assessed through mutagenesis of the *Sox2* locus in ES cells (Huang et al., 2021), as well as their cooperative and redundant functions when disposed as arrays at the *Pax3* locus (Anania et al., 2021). In this study, we genetically dissected a series of five CBSs all located on one side of the strong TAD boundary located within the *HoxD* gene cluster. In addition to being involved in the making of a tight chromatin border (Rodriguez-Carballo et al., 2017), these sites delimit various subsets of contiguous *Hoxd* genes, which respond to distinct tissue-specific remote enhancers thus raising the possibility that CBS-dependent micro-chromatin domains could be defined for each (series of-) enhancer(s).

### CTCF as moderator of spatial colinearity

We used our allelic series of mutations *in cis* to try to distinguish between a micro-insulating effect (between genes), a macro-insulating effect (between TADs) or an anchoring function for these multiple CBSs. The efficiencies of the mutations were checked using both CTCF and RAD21 in ChIPmentation experiments, which revealed that CTCF binding had been fully abrogated and that RAD21 was no longer enriched at these sites. No cryptic CBS was revealed even in the absence of the five native CBSs and the enrichment of RAD21 was redistributed towards the remaining CBSs, with a clear preference for the CBSs located close to the deleted series (e.g. CBS4 in the Del(CBS1-3) allele), suggesting that cohesin-driven chromatin architectures were locally modified in these mutants, at least by using a mix of trunk cells as starting material. However, CBS5 displayed only a moderate increase in RAD21 signal upon deletion of its four 3’ neighboring CTCF, suggesting that sequence- or context-dependent factors, others than motif orientation, can influence the capacity of CBS to retain cohesin (Phillips-Cremins et al., 2013).

This observation was paralleled by a more condensed aspect of the cluster itself, as determined by chromosome capture, with intra-cluster interactions increasing along with the number of CTCF sites mutated and progressively extending towards the centromeric end of the gene cluster up to the TAD boundary in the full CTCF mutant. Concomitantly, long-range contacts between the cluster and T-DOM were lost, an effect particularly visible at positions corresponding to the presence of convergent CTCF sites such as the CS38-40 region as well as other CBSs located further telomeric. We interpret these two phenomena as a single effect of removing all sites orientated towards T-DOM within the *Hox* cluster; the lack of long-range interactions with T-DOM somehow relaxed the architecture, thus allowing *Hoxd* genes to establish local interactions, which are normally reduced whenever the cluster in under tension through its contacts with T-DOM.

How such an increase in local interactions may affect *Hox* gene transcription is more difficult to evaluate in our experimental paradigm. *Hox* genes are transcribed in successively more ‘posterior’ combinations along with embryonic caudal trunk extension, with a correspondence between the gene’s order in the clusters and the anterior to posterior (A-P) level where they become activated (Gaunt et al., 1988). Previous deletions on CTCF sites showed that this particular A-P level could be modified in some instances, leading to a local mis-expression of neighbouring genes (Narendra et al., 2016a). In this view, particular CBSs could be considered as having an insulator function. Here, by using this complete allelic series, we show that this effect is systematically observed for genes located close to the deleted CBSs. However, while the positioning of the expression boundaries was abnormal in the multiple mutated Del(CBS1-4) and Del(CBS1-5) specimens, the colinear distribution of these boundaries was conserved, suggesting that while CTCF is required to fine-tune the A-P levels where particular genes will be switched on, it is not necessary for the implementation of spatial colinearity in itself. The proper adjustment of these expression domains is required for harmonious development, as shown by the phenotypic alterations accompanying these allelic series, which were all in agreement with previous results and general principles of homeosis (Gerard et al., 1996; Kessel and Gruss, 1991b; Narendra et al., 2015; Tarchini et al., 2005). Therefore, in this context, CTCF can be seen as a moderator of spatial colinearity rather than its organizer.

### CTCF for anchoring enhancers or for insulating their effects?

In the case of remote enhancers-promoter interactions, bound CTCFs were shown to act both as insulators and as anchoring elements. For example, a tandem insertion of CTCF sites could insulate the *Pcdh* promoter from an enhancer (Jia et al., 2020) and tissue-specific insulation of a CTCF bound region was also reported to control the selective expression of human growth hormone hGH) gene cluster (in the placenta (Tsai et al., 2016). In contrast, other CTCF bound regions can act as facilitators/ tethering elements to bring enhancers located at long distances close to their promoters, as exemplified by the transcription of the *Shh* gene during limb development, which requires the proximity of the ZRS sequence, which is achieved either partly (Paliou et al., 2019) or fully (Ushiki et al., 2021) by the presence of CTCF sites.

Our mutant alleles revealed that, while CBSs were generally used as insulators, the insulating function was in some cases tissue-specific, for the observed extension of the expression domain was not the same in all tissues analysed. Some CBSs were also necessary to properly anchor enhancer-promoter interactions, as reported in other loci, such as the Thy1 locus (Ren et al., 2017). For example, the clear gain of expression of *Hoxd9* in the FMPs observed in both Del(CBS1-2) and Del(CBS1-3) was reduced or mostly abrogated in the Del(CBS1-4) and Del(CBS1-5) alleles, respectively, suggesting that both CBS4 and CBS5 triggered the appropriate contact rather than insulating yet another transcription unit from remote enhancers. This hypothesis is further supported by the observation that although *Hoxd11* is only weakly upregulated in the FMPs of Del(CBS1-4) embryos, correlating with the weak RAD21 enrichment of CBS5 in these mutants, its gain was completely abolished in Del(CBS1-5) homozygous mutant specimen. In fact, the anchoring and insulating functions could be exerted by the same site in different contexts, as shown with CBS2, which clearly acted as an anchor in mammary bud cells where the expression of *Hoxd9* was lost when CBS2 was mutated, whereas the same gene was upregulated in both paraxial and lateral mesoderm of CBS2 mutant embryos. Such a dual role has been reported for other CTCF binding sites such as those flanking the HS5 enhancer of *Protocadherin a* cluster (Guo et al., 2015, 2012; Jia et al., 2020), and the insulator activity of CTCF bound elements has been related with their CpG methylation and epigenetic modifiers state in different contexts (Bell and Felsenfeld, 2000; Guo et al., 2012).

The functional ambivalence of these CBSs may depend on several parameters such as the presence of other tissue-specific factors, the local extrusion conditions or the strength of interactions, to mention a few possibilities. In any case, the dual and context-dependent activity of CBSs we report at this locus indicate that one should be careful when assessing the effects of mutating one or a few CTCF sites within a larger array of such sequences, for the overall effect of such arrays may not be extrapolated from the addition of single mutation events. The functional compensation we observed when mutating the full series of CBSs (Del(CBS1-5) illustrates this point.

### CBSs in the making of the TAD boundary at the *HoxD* cluster

All four mammalian *Hox* clusters are partitioned in between two TADs, with a boundary systematically isolating the most posterior group 13 *Hox* gene from the others. Group 13 *Hox* genes are coding for proteins that participate to the morphogenesis of terminal structures and hence embryo development must be protected from the action of such proteins, which can abrogate axial extension (Young et al., 2009) or affect proper morphogenesis (Bolt et al., 2021; Darbellay et al., 2019) through their dominant negative function over other HOX proteins. This isolation is achieved through the positioning of *Hox13* genes into a different TAD, thus making them unresponsive to the numerous enhancer sequences acting upon other *Hox* genes during development. As illustrated with *HoxD*, TAD boundaries at *Hox* cluster are remarkably tight (Rodriguez-Carballo et al., 2017). While, as for many other such chromatin borders, *Hox* loci display arrays of CBSs (Gómez-Marín et al., 2015; Kentepozidou et al., 2020; Vietri Rudan et al., 2015), the orientations of these sites are generally not as intermingled as is often observed (e.g. (Anania et al., 2021; Nanni et al., 2020)). Instead, the CBSs are organized as two arrays of opposite orientations thus defining a clear point of transition, tightly isolating *Hox13* from the gene cluster.

While several studies have revealed that the number of CBS was important for insulation at TAD boundaries, likely through a synergistic effect (Huang et al., 2021), particular CBS may be more important than others in achieving this task (Anania et al., 2021) and it appears that tandem distributions of CBS may provide robustness to the TAD border. In our allelic series, the global insulation capacity of the TAD border was increasingly affected along with more CTCF sites being mutated, as best seen in the case of the developing limbs. However, this effect was weak and the TADs remained almost as in control cells, even when all CBSs with the telomeric orientation had been mutated. Strong variations were observed in the interactions between the cluster and T-DOM, yet the TAD border itself was only weakly affected, suggesting that a single array of CTCF sites with the same (centromeric) orientation is sufficient to provide isolation between the TADs. This was confirmed by the lack of observable gain of expression of *Hoxd13* whenever it was tested, in particular in proximal limb cells, and this even though the position of the TAD border was shifted towards this gene. This illustrates that the strong T-DOM enhancers, actively working on *Hoxd10* and *Hoxd11*, were still unable to reach *Hoxd13*, even in mutant Del(CBS1-5) limbs and hence it suggests that a single array of CTCF sites with the same orientation is sufficient to reach a fair level of insulation, at least in this context. However, the shift of the TAD border towards *Hoxd13* may clearly weaken the isolation of this gene and we cannot rule out its ectopic transcription in response to T-DOM enhancers in a particular cellular context, during development or adulthood. Therefore, the deleted CTCF sites may function to tighten the TAD border, rather than to establish it.

These multiple functions for such CBSs arrays at *Hox* loci may explain the remarkable evolutionary conservation in the global organisation of such binding sites. Indeed, despite several hundred millions years of separate evolution for these paralogous gene clusters (Holland et al., 1994) accompanied by drastic functional divergences (Soshnikova et al., 2013) and sometimes the loss of several genes (Hoegg and Meyer, 2005), the general distribution of two these arrays of CTCF sites, as well as their opposed orientation were conserved. Likewise, human and mouse *Hox* clusters are identical in this respect, unlike other situations where the function of CTCF sites was conserved, but not their exact locations (Ushiki et al., 2021). Furthermore, the fact that in all four clusters the first CTCF site is located right upstream of the group 4 *Hox* gene promoter demonstrates a function for these CBSs more precise and complex than making a mere TAD border.

### The impact of CBS for *Hoxd* gene regulation in limb buds

During limb development, various subsets of *Hoxd* genes respond first to T-DOM-located proximal enhancers, then to C-DOM-located digit enhancers (Andrey et al., 2013). Because the two subgroups of target genes are delimited by CTCF sites, it was suggested that such CBSs may specify, through a 3D structure, particular enhancer-promoter interactions leading to these bimodal transcription patterns (Rodríguez-Carballo et al., 2020). However, the most visible effect observed across the different mutant lines generated in this study was more quantitative than qualitative and no obvious mis-expression of target gene subgroups was scored.

Furthermore, all variations observed in the amounts of mRNAs in all mutant configurations tested were systematically losses of transcripts, when compared to control mRNA levels. Such losses could be explained when looking at variations in chromatin topology. For example, mutations in all five CTCF sites considerably weakened contacts between the cluster and T-DOMb, where most proximal enhancers are located (Andrey et al., 2013; Rodríguez-Carballo et al., 2020). These contacts were redirected towards T- DOMa, which is poor in proximal enhancers, but rich in other regulations, leading to the ectopic expressions described above, concomitantly to a loss of expression in proximal limbs.

While the overall weakening of contacts in T-DOMb was even more evident in Del(CBS1-5) mutants, this abrogation of all five CTCF sites resulted in a re-equilibration of contacts with T-DOMa. In addition, the increase in local intra-TAD contacts observed in the C-DOM of Del(CBS1-3) mutants was also reverted in the distal forelimb of Del(CBS1-5) mutant embryos, likely accounting for the rescue in *Hoxd* gene expression levels observed in these mutants. These results indicate that CBS1 to 5 are required for the correct anchoring of *Hoxd* genes with their long-range enhancers situated in the flanking TADs and the mutagenesis of some of these sites thus impacts upon their interaction pattern, as well as on the internal TAD architecture. However, the removal of all sites resulted in a more homogeneous loss of contacts between *Hoxd* genes and their regulatory landscapes. Therefore, in this context at least, rather than being used to fine select the appropriate sets of target *Hoxd* genes, the CTCF sites are required to secure the highest possible expression levels by triggering more robust interactions with T-DOMb-located long-range enhancers. In this view, CBSs may achieve a function in potentiating regulations rather than in their specific establishment.

### A gene cluster under tension

How such a potentiation effect could be achieved is elusive and the abrogation of every single CTCF binding site taking part in these regulations might be of help in this context. It is likely that this effect results from a global re-organisation of the chromatin landscape rather than to punctual modifications. For example, in the trunk of Del(CBS1-5) mutant specimen, anterior *Hoxd* genes (i.e., *Hoxd3*, *Hoxd4*) established ectopic interactions with more ‘posterior’ genes (*Hoxd9*, *Hoxd10*). Concomitantly, the ‘anterior part’ of the gene cluster loses many interactions with T-DOM, suggesting that the two patterns of interactions are exclusive and illustrate two global conformations of the 3D chromatin at the locus. As a consequence, it is possible that, in the normal situation, the contacts established with T-DOM are necessary to bring the gene cluster under some tension, to open it such that it may respond to enhancers in an optimal manner. In this view, CTCF may be considered as a factor used to maximize transcription through the re- organization of chromatin, rather than a key determinant for enhancer-promoter interaction, necessary for transcription to occur. Our observations are in agreement with the somehow reduced effect of removing CTCF genome wide upon global transcription patterns, at least during development (Kubo et al., 2021; Nora et al., 2017; Soshnikova et al., 2010; Wan et al., 2008)

## ETHICS APPROVAL

All experiments involving animals were performed in agreement with the Swiss Law on Animal Protection (LPA), under license No. GE 81/14 (to DD).

## DATA AVAILABILITY

All raw and processed datasets are available in the Gene Expression Omnibus (GEO) repository under accession number GSE181387. All scripts necessary to reproduce figures from raw data are available at https://github.com/lldelisle/scriptsForAmandioEtAl2021.

## AUTHORS CONTRIBUTIONS

R.A.: Designed and conducted experiments, analyzed datasets, formalized results and wrote the paper. L.B.: Designed and conducted experiments, analyzed datasets, formalized results and wrote the paper. L.L.-D. : Analyzed and evaluated the statistical significance of datasets and wote the paper.

B.M. : Designed, produced, genotyped and helped analyze mouse mutants.

J.Z.: Analyzed and evaluated the phenotypes of mutant animals.

S.G. : Genotyped mouse mutants and helped for mouse work.

D.D. : Designed experiments, transported mice and wrote the paper.

## ACKNOWLEDGEMENTS

We thank Aurélie Hintermann for communicating results prior to publication, Dr. Chase Bolt for providing early access to control cHi-C datasets, Dr. Hocine Rekaik for help in ChIP-M experiments, as well as other members of the Duboule laboratories for comments and discussion. We are grateful to Thi Hanh Nguyen Huynh for her help with mice breeding and genotyping. We thank the iGE3 Genomics Platform of the University of Geneva and the Gene Expression Core Facility at the School of Life Sciences of EPFL for their assistance in NGS related experiments. We are grateful to the small animal preclinical imaging platform of the Medical Faculty of Geneva for micro-CT scans imaging. Part of the calculations have been performed using the facilities of the Scientific IT and Application Support Center of EPFL.

## MATERIALS AND METHODS

### Cloning of sgRNAs and generation of the CBS mutant stocks

The sgRNA targeting guides were generated by annealing complementary pairs of oligonucleotides (Supplemental Table S2) and cloning into the pX330 vector as described in (Darbellay et al., 2019). For sgRNA transcription, we PCR amplified the sgRNA sequence cloned into the px330 plasmid using a T7 promoter containing primer and a universal reverse oligonucleotide (TAATACGACTCACTATAG). PCR products were gel purified and transcribed *in vitro* using the HiScribe™ T7 High Yield RNA Synthesis Kit (NEB). Cas9 mRNA was synthesized from the pX330 plasmid using the mMESSAGE mMACHINE® T7 Ultra (Thermofisher) according to manufacturer instructions. The transcribed sgRNAs were purified using the RNeasy mini kit (Qiagen). The purified Cas9 mRNA and sgRNAs were co-electroporated in mouse fertilized oocytes. The list of sgRNAs used is given in Table S2.

With the exception of the Del(CBS1-4) line, F0 animals obtained upon Cas9/sgRNA electroporation were screened at weaning age using the Surveyor Mutation Detection Kit and specific primers amplifying a fragment 500bp to 1000bp large around the targeted CBS (Supplemental Table S3). The Del(CBS1-4) animals were screened by PCR (Supplemental Table S3). The first mutant strain obtained was the Del(CBS2). The Del(CBS1-2) was obtained on top of Del(CBS2) zygotes by introducing the CBS1’ mutation depicted in Fig. S4. The distinctive CBS1 mutation (Supplemental Fig. S4) was present only in the Del(CBS1) strain, produced independently. The Del(CBS1-3) strain was produced on top of Del(CBS1-2) zygotes and the Del(CBS1-4) strain was produced on top of Del(CBS1-3) zygotes, with the mutation referred to as CBS4 in Supplemental Fig. S4. The Del(CBS1-5) strain was produced on top of another Del(CBS1-4) allele containing the CBS4’ mutation referred to in Fig. S4.

For genotyping, specific primers were used to amplify a region of ca. 160bp to 423bp surrounding the newly mutated CBS for each combined mutant strain generated, allowing to discriminate the WT from deleted alleles in 5 % agarose gel electrophoresis (Supplemental Table S3). All mutant alleles were verified by sanger sequencing (Supplemental Fig. S4). To evaluate whether any new mutated CBS was positioned *in-cis* or *in-trans* with previous CBS mutation(s), several F0 specimen for the new allele were crossed over WT animals and F1 progenies genotyped to look at the segregation of mutations. In this way, mutant alleles were recovered with the newly mutated CBS positioned either *in cis* or *in trans* with previous CBS mutations. All mice used either for zygotes electroporation or for further breeding were from a mix Bl6XCBA background, as all *HoxD* alleles produced in the laboratory.

### Whole-mount *in situ* hybridization (WISH)

WISH experiments were performed as described in (Woltering et al., 2009). The *Hoxd11*, *Hoxd10*, *Hoxd9*, *Hoxd8*, *Hoxd4* gene probes were described in (Dolle et al., 1991; Gerard et al., 1996).

### Micro-CT scan analyses

Mouse adult skeletons were revealed after micro-CT scanning using the Quantum GX2 micro-CT Imaging System (PerkinElmer) at the small animal preclinical imaging platform of the Medical Faculty of the University of Geneva. Images were analyzed using the OsiriX MD v.10.0.1software.

### ChIPmentation (ChIP-M)

ChIP-M experiments for CTCF and RAD21 were performed according to (Darbellay et al., 2019). Briefly, trunk regions of E10.5 embryos (from below the post-occipital region) derived from trans-heterozygous crosses were individually dissociated into single cells with collagenase and fixed in 1% formaldehyde solution for 10 minutes at room temperature. The crosslinking reaction was stopped adding Glycine to a final concentration of 0,125 M and the cell pellet was washed three times with cold PBS with protease inhibitors (Complete mini EDTA –free proteinase inhibitor cocktail; Roche). Fixed samples were stored at -80C. Yolk sacs were used for genotyping. Samples from homozygous mutant embryos or control littermates were resuspended in sonication buffer (Tris HCl pH=8.0 50mM; EDTA 10 mM SDS 0,25%; protease inhibitors) and the chromatin was sheared in a Covaris S200 sonicator to an average fragment size of 250-300 bp. The sonicated chromatin was diluted 2.5 times with dilution buffer (HEPES pH=7.3 20mM; EDTA 1mM; NP40 1%; NaCL 150mM; protease inhibitors). Antibodies against CTCF (Active Motif #61311; 4gr) or RAD21 (Abcam # ab992; 5ugr) were conjugated with Dynabeads Protein G (Thermofisher) magnetic beads. The bead-antibodies complexes were added to the diluted chromatin and incubated overnight at 4°C in a rotating platform. The day after, chromatin-antibody complexes were washed using a magnetic stand with RIPA buffer (Tris HCl pH=8.0 10mM; EDTA 1mM; Sodium Deoxycholate 0,1% TritonX-100 1%; NaCl 140mM; two washes); RIPA-High salt buffer (Tris HCl pH=8.0 10mM; EDTA 1mM; Sodium Deoxycholate 0,1% TritonX-100 1%; NaCl 500mM; two washes), LiCl-buffer; (Tris HCl pH=8.0 10mM; EDTA 1mM; LiCl 250mM; Sodium Deoxycholate 0,5% NP40 0,5%; two washes) and Tris HCl 10mm pH=8.0 (two washes). The chromatin was then tagmented for two minutes with the Tn5 transposase (Illumina). Tagmented chromatin was eluted, reverse crosslinked and purified using Qiagen Minielute columns. Libraries were quantified using the Kapa library quantification kit and PCR amplified using barcoded primers. ChIP-M libraries were pair-end sequenced in an NextSeq 500 sequencer (PE 2x 37-43bp).

Adapters and bad quality bases were removed from the fastqs with cutadapt version 1.16 (Martin, 2011) (- a CTGTCTCTTATACACATCTCCGAGCCCACGAGAC-A <u>CTGTCTCTTATACACATCTGACGCTGCCGACGA -q 30 -m 15)</u>. Filtered reads were mapped to the mouse genome mm10 with bowtie2 version 2.3.5 (Langmead and Salzberg, 2012) with default options). Only alignments with a mapping quality above 30 were kept (Danecek et al., 2021). PCR duplicates were removed with Picard version 2.19.0 (Broad Institute, n.d.). To decrease the importance of fragment length variation between libraries only the first read in pairs were kept and BAM was converted to BED. Peak calling was run with a fixed fragment size of 200bp with macs2 callpeak version 2.1.1.20160309 ( --call-summits -f BED --nomodel --extsize 200 -B --keep-dup all). In order to normalize all ChIP-M despite different signal to noise ratios, we run MAnorm version 1.1.4 (Shao et al., 2012) with -w 100) between each sample and the second replicate of the wildtype. The M-A model coefficient was extracted and used to normalize each bedgraph from macs2 (see https://github.com/lldelisle/scriptsForAmandioEtAl2021 for all details). Replicates were then averaged at each base using bedtools version 2.27.1 (Quinlan, 2014). In supplemental Figure S9, the coverages from macs2 were normalized to the million tags. Then wildtype replicates were averaged.CTCF site orientation was determined using CTCFBSDB 2.0 (http://insulatordb.uthsc.edu/) (Ziebarth et al., 2012) using the CTCFBS Prediction tool with sequences of 500bp centered on each summit of each of the three wildtype replicates. For each sequence, the motif with the highest score among REN_20, LM2, LM7 and LM23 motifs was kept. Only motifs that are common between the three replicates are displayed.RAD21 MAnorm-normalized coverage were quantified on 500bp regions centered on the CTCF motifs with multiBigwigSmmary from deepTools version 3.5 (Ramirez et al., 2016). The PWM motif logos of Supplemental Fig. S4 were generated using the STAMP (Mahony et al., 2007; Mahony and Benos, 2007).

### ChIP-seq

CTCF ChIP-seq experiments in supplemental Figure S1 were performed as described in (Rodriguez-Carballo et al., 2017). For the posterior trunk samples, the portion corresponding to the tailbud, presomitic mesoderm and 2 to 3 somite pairs, excluding as much as possible the region of the incipient hindlimb buds, was micro-dissected from approximately thirty E9.75 to E10 embryos. For the brain samples, a pool of six micro-dissected forebrains from E12.5 embryos was used. The ChIP-seq fastqs of E12.5 distal and proximal forelimb samples were downloaded from GEO (GSM2713707 and GSM2713708, respectively). TrueSeq adapters were removed from single-reads fastqs with cutadapt version 1.16 (Martin, 2011) <u>-a CTGTCTCTTATACACATCTCCGAGCCCACGAGAC -q 30 -m 15)</u>. Filtered reads were mapped to the mouse genome mm10 with bowtie2 version 2.3.5 (Langmead and Salzberg, 2012) with default options). Only alignments with a mapping quality above 30 were kept (samtools version 1.9 (Danecek et al., 2021). Peak calling was run with a fixed fragment size of 200bp with macs2 callpeak version 2.1.1.20160309 (-- call-summits --nomodel --extsize 200 -B). Finally, the coverages were normalized to the million unique tags.

For the published ChIP-seq used in supplemental Figure S3, coverage files were downloaded from GEO as well as the corresponding peaks files. To identify CTCF orientation, the procedure was very similar to the analysis for ChIP-M except that for the human there was no summit information so the peaks were scaled to 500bp.

### Capture Hi-C

Micro-dissected E12.5 proximal and distal forelimb pairs and individual E9.5 trunks were isolated in PBS supplemented with 10% Fetal Calf Serum, dissociated into single cell by collagenase treatment, fixed in 1% formaldehyde and stored at -80°C until further processing. Samples were genotyped by PCR as described above to select for homozygous mutant or control tissues. The SureSelectXT RNA probe design and Capture Hi-C experiments were performed as described in (Bolt et al., 2021). The first part of the data analysis was performed on a local galaxy server (Afgan et al., 2016). Raw reads were preprocessed with cutadapt version 1.16 (Martin, 2011) (-a AGATCGGAAGAGCACACGTCTGAACTCCAGTCAC - A AGATCGGAAGAGCGTCGTGTAGGGAAAGAGTGTAGATCTCGGTGGTCGCCGTATCATT --minimum-length=15 --pair-filter=any --quality-cutoff=30). Then hicup version 0.6.1 (Dryden et al., 2014; Langmead and Salzberg, 2012) and samtools 1.2 (Danecek et al., 2021) were used with default parameters. The bam file was converted to tabular file with a python script (see https://github.com/lldelisle/scriptsForAmandioEtAl2021 for more details). The pairs were then loaded to a 10kb resolution matrices with cooler version 0.7.4 (https://doi.org/10.1093/bioinformatics/btz540). The heatmaps in figures 1, 3, 4, 5 and Supplemental Figures S6, S7 and S9 are representations of the two replicates using HiCExplorer hicSumMatrices tool version 3.6 (https://doi.org/10.1038/s41467-017-02525-w https://doi.org/10.1093/nar/gky504 https://doi.org/10.1093/nar/gkaa220) and cooler balance. Subtraction maps were generated from balanced matrices using the HiCExplorer hicCompareMatrices tool version 3.6. For Figure S6, a correction was applied to account for differences between matrices in their close distance-dependent signal, using the HiCExplorer hicTransform tool version 3.6 (Ramírez et al., 2018; Wolff et al., 2020) with the obs_exp_non_zero transformation method and the perChromosome option. The TAD separation scores were computed with HiCExplorer hicFindTADs version 3.6. (Ramírez et al., 2018; Wolff et al., 2020) with a fixed window size of 240kb. Heatmaps were plotted with pyGenomeTracks 3.5 (Lopez-Delisle et al., 2021; Ramírez et al., 2018) Virtual C profiles were generated similarly to (Despang et al., 2019), with a custom python script available on https://github.com/lldelisle/scriptsForAmandioEtAl2021. The viewpoints coordinates (mm10) used are: *Hoxd4*, chr2:74721977; *Hoxd8*, chr2:74704614; *Hoxd9*, chr2:74697726; TAD border: chr2:75588758. The quantifications in Figures S6, S7 and S13 were also carried out with a custom python script.

### RNA-seq

Micro-dissected individual pairs of either control or mutant E12.5 proximal and distal forelimbs were stored at -80°C in RNAlater stabilization reagent (Ambion) before further sample processing. Total RNA was extracted from tissues using Qiagen RNeasy Plus Micro Kit (Qiagen) after disruption and homogenization according to manufacture instructions. RNA quality was assessed using an Agilent 2100 Bioanalyser. The sequencing libraries were prepared according to TruSeq Stranded mRNA Illumina protocol, with polyA selection. RNA-seq libraries were sequenced on an Illumina HiSeq 4000 sequencer, as single-reads (read length 50 bp). Raw RNA-seq reads were processed with cutadapt version 1.16 (Martin, 2011) <u>-a CTGTCTCTTATACACATCTCCGAGCCCACGAGAC -q 30 -m 15)</u> to remove TruSeq adapters and bad quality bases. Filtered reads were mapped on the mouse genome mm10 with STAR version 2.7.0.e (Dobin et al., 2013) with ENCODE parameters with a custom gtf file (https://doi.org/10.5281/zenodo.4596489) based on Ensembl version 102. FPKM values were evaluated by cufflinks version 2.2.1 (Roberts et al., 2011; Trapnell et al., 2010) with options --max-bundle-length 10000000 --multi-read-correct --library-type “fr- firststrand” -b mm10.fa --no-effective-length-correction -M MTmouse.gtf -G). Counts from autosomal chromosomes were used for differential expression analysis with DESeq2 version 1.24.0 (Love et al., 2014) with R version 3.6.0 (www.r-project.org) with default parameters except for theta, which was fixed to 0.15 and 0.99. Only genes with absolute log2 fold-change above 0.58 and adjusted p-value below 0.05 were considered as significant. All the significant results are summarized in Supplemental File 1. PCA and clustering were performed on log2(1 + FPKM) values of the 500 most variant genes from autosomal chromosomes.

## LEGENDS TO SUPPLEMENTAL FIGURES

**Supplemental Figure S1.**
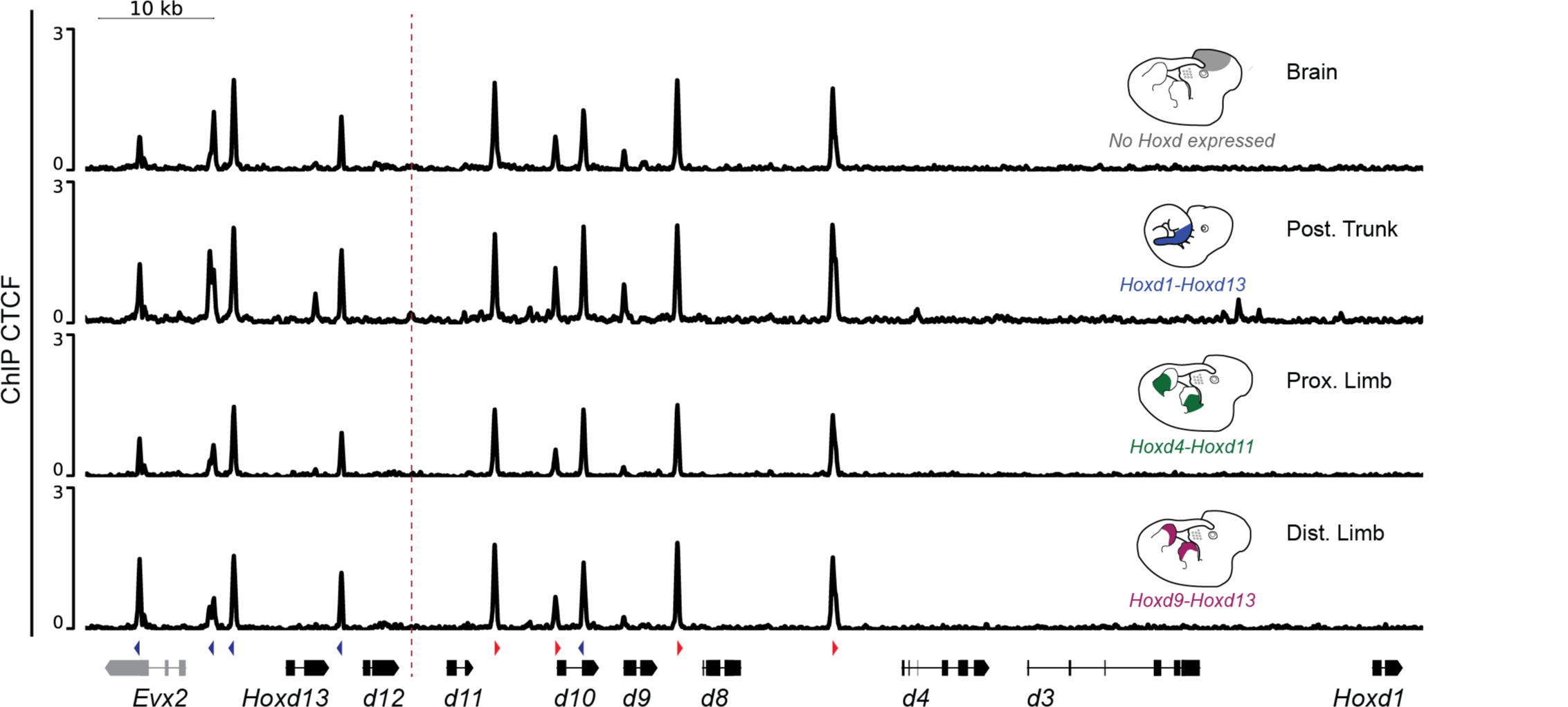
Constitutive CTCF binding at the *HoxD* locus. CTCF ChIP-seq profiles over the *HoxD* cluster (mm10: chr2:74650810-74767377) in different mouse embryonic tissues as indicated in each panel. Schematics on the right represent the dissected samples and indicate the genes expressed in these samples. ChIP-seq datasets from E12.5 proximal and distal limbs (two bottom tracks) are from (Rodriguez-Carballo et al., 2017) (GSM2713708 and GSM2713707, respectively). The orientations of the CBSs are indicated with arrowheads below the bottom track, with red for a telomeric orientation (pointing towards T-DOM) and blue for a centromeric orientation (pointing towards C-DOM). *Hoxd* genes and *Evx2* are depicted in black and grey, respectively. Red dashed line indicates change in orientations of CBS.

**Supplemental Figure S2.**
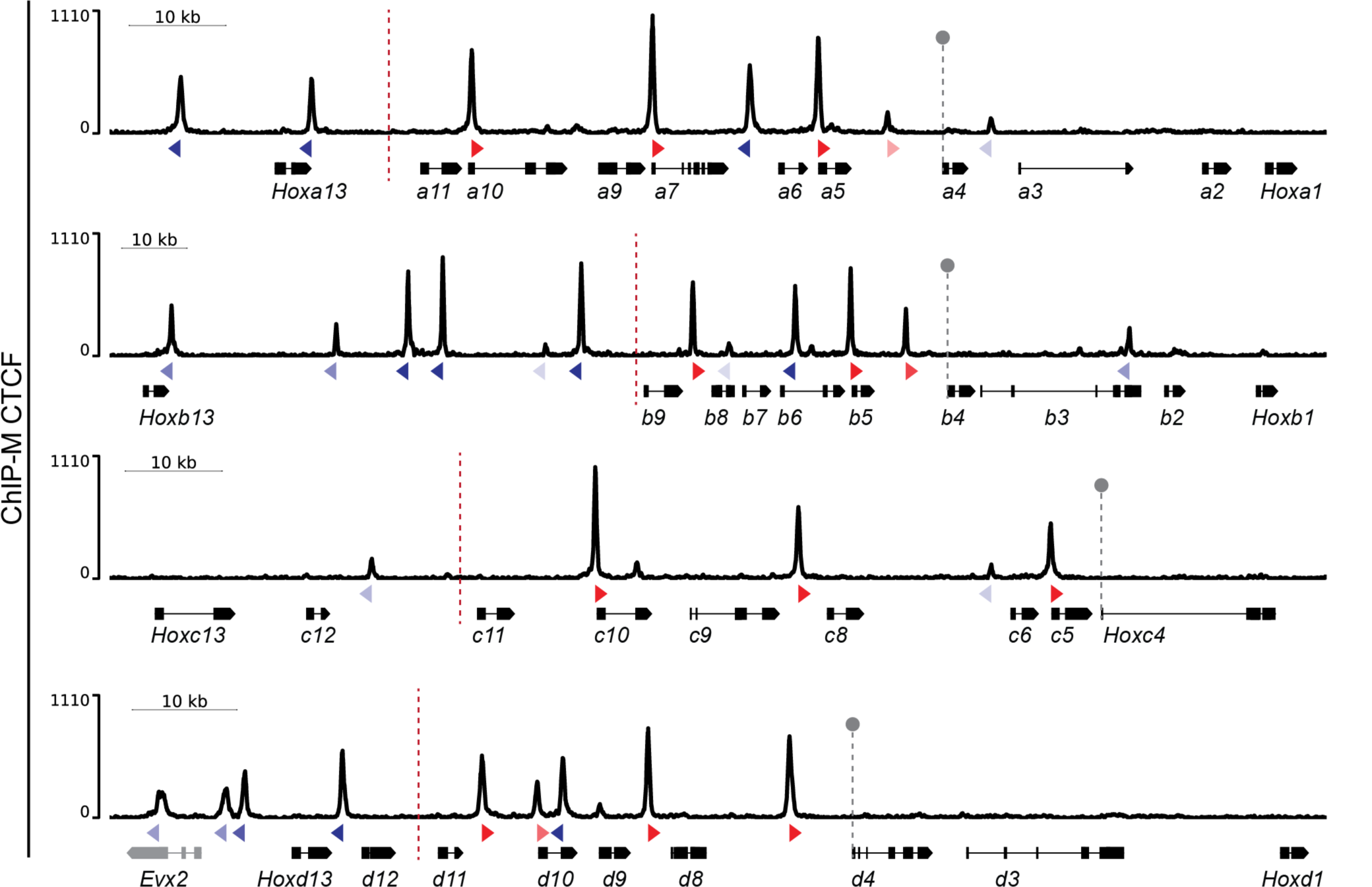
CTCF binding profiles at paralogous *Hox* clusters. CTCF ChIP-M profiles using E10.5 trunk cells over the *HoxA* (mm10: chr6:52151994-52278047), *HoxB* (mm10: chr11:96189088- 96376579), *HoxC* (mm10: chr15:102916367-103042207) and *HoxD* (mm10: chr2:74650810-74767377) paralogous clusters. The orientations of CBSs are shown as in Fig. S1. Colour intensity represents scale of peak coverage (darker colour higher, lighter colour lower). The *HoxA* cluster is depicted in the reverse genomic orientation for the ease of comparison. The dashed red line represents the region of inversion in CBS orientation, matching TAD boundaries. The grey pin line points to the promoter of *Hox* group 4 genes, located near the first CBS in all four cases.

**Supplemental Figure S3.**
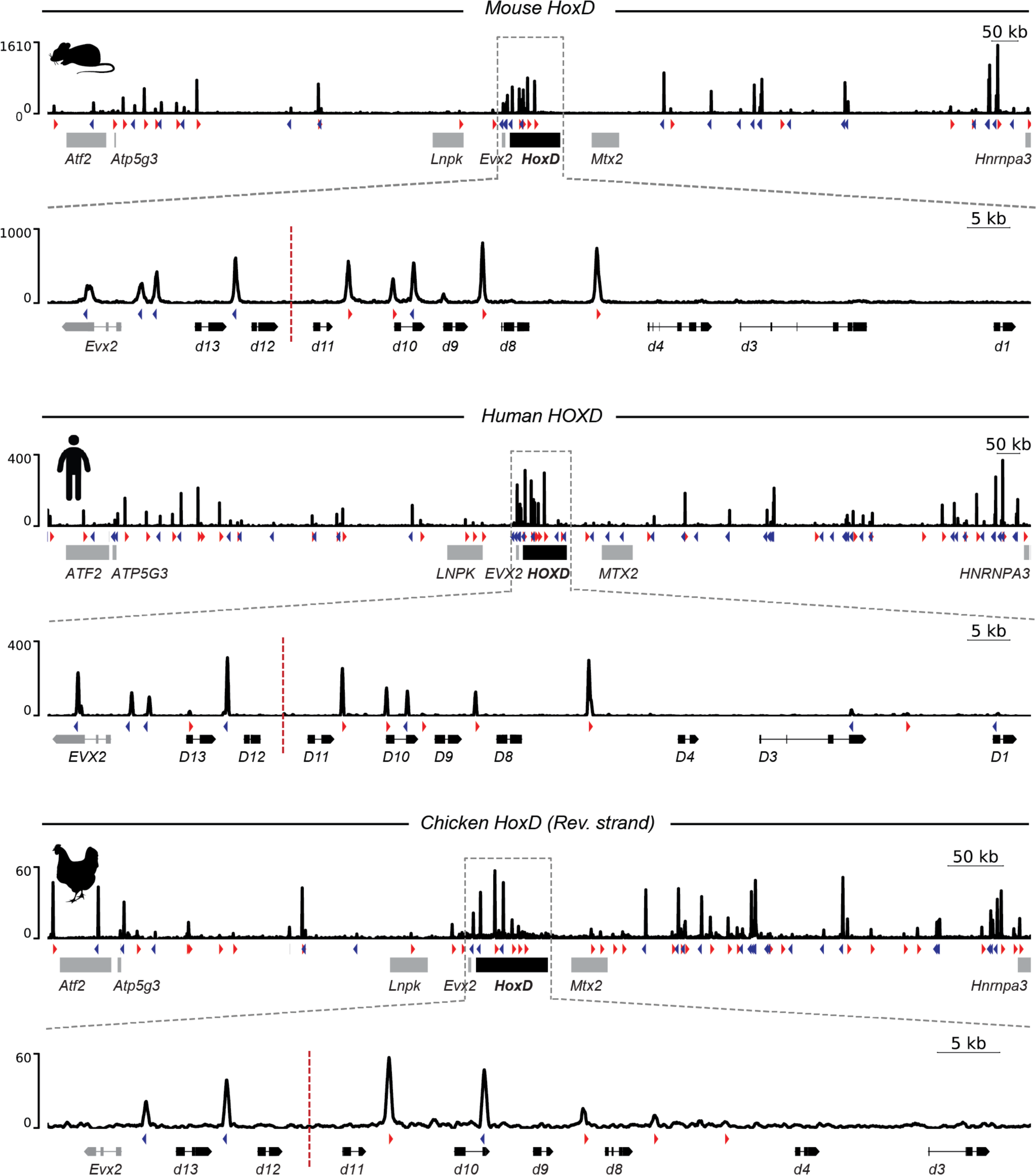
Evolutionary conservation of CBSs at the *HoxD*. CTCF ChIP-M or ChIP-seq profiles over the human, mouse and chick *HoxD* locus. Mouse ChIP-M profiles are from E10.5 trunk cells (mm10: chr2:73779626-75669724 and chr2:74650810-74767377), whereas the ChIP-seq dataset is from human H1 ES cells (ENCODE Project Consortium, 2012) (hg19: chr2:175895200-178092461 and chr2:176941129-177057814) and the chick ChIP-seq data from HH20 forelimbs (Yakushiji-Kaminatsui et al., 2018)(GSM3182452)(galGal5: chr7:15889800-16780000 and chr7:16323377-16402356). For each species, CTCF binding profiles are shown for both the genomic regions spanning the two TADs (Top) and the *HoxD* cluster (bottom). The chicken *HoxD* genomic region is represented in the reverse genomic orientation for the ease of comparison. The orientations of CBSs are as for Fig. S1.

**Supplemental Figure S4.**
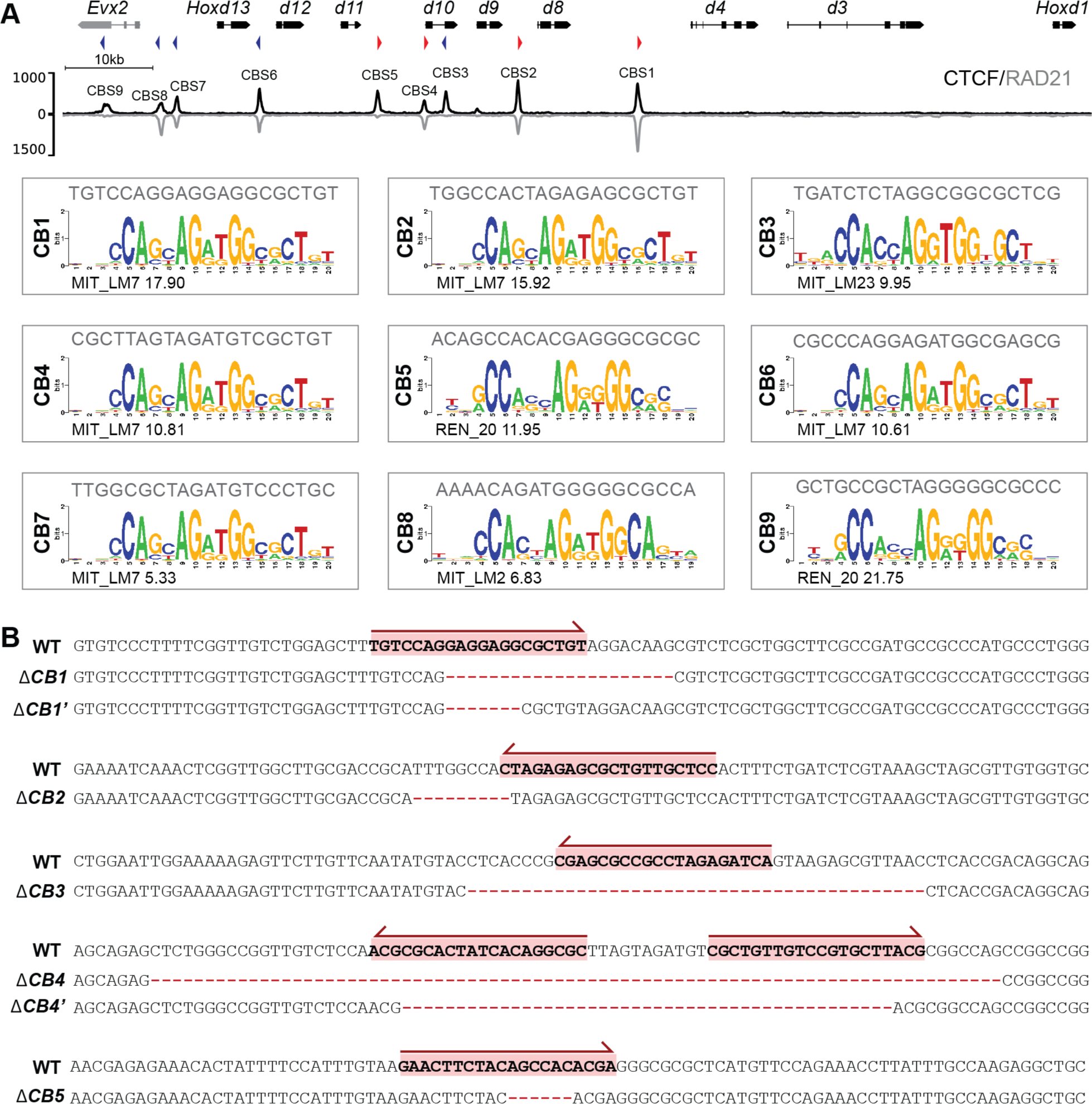
CRISPR-Cas9 mediated CBS mutagenesis. **(A) Top.** CTCF (black) and RAD21 (grey) ChIP-M profiles of E10.5 embryonic trunk cells covering the *HoxD* cluster (mm10): chr2:74650810-74767377). CBSs are indicated above the peaks with their orientations atop. **Bottom**. CBS predictions were performed using the CTCFBSDB 2.0 tool (Ziebarth et al., 2012) see Material and Methods). For each CBS, the CTCF position weight matrix (PWM) giving the highest prediction score (indicated below) is shown. On top of the PWM the motif sequence identified is represented (**B)** Sequence alignments showing the wildtype version of the different CBSs mutagenized in this study and their respective deletion alleles (red dashes represent deleted nucleotide positions). The spanning of the Cas9 sgRNA is in red. Red arrows indicate the sgRNA strand specificity. For CBS4, two sgRNA guides were electroporated in fertilized mouse oocytes. The 1CBS1 sequence corresponds to the CBS1 deletion of the Del(CBS1) allele, whereas the 1CBS1’ alignment corresponds to the CBS1 deletion generated in the Del(CBS1-2), Del(CBS1-3), Del(CBS1-4) and Del(CBS1-5) alleles. The 1CBS4 sequence corresponds to the CBS4 deletion in the Del(CBS1-4) allele, while 1CBS4’ indicates the sequence present in the Del(CBS1-5) allele.

**Supplemental Figure S5.**
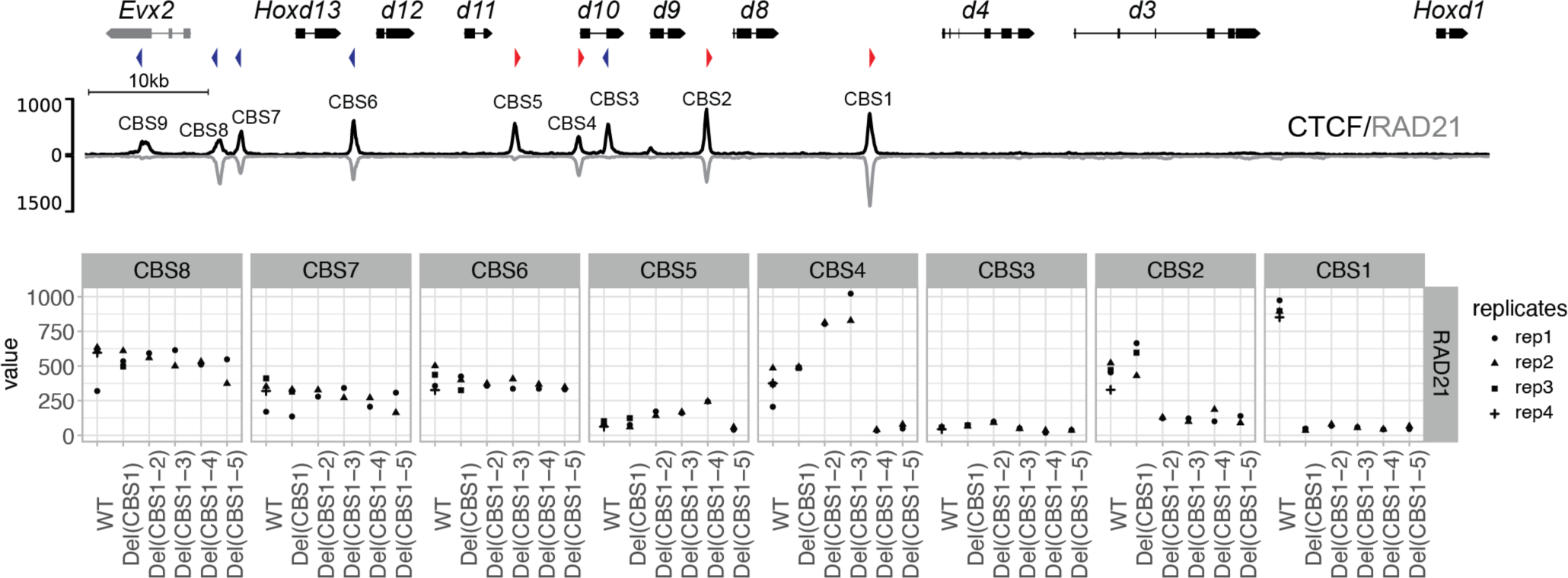
Quantification of RAD21 at *HoxD* CBSs. **Top.** CTCF (black) and RAD21 (grey) ChIP-M profiles of E10.5 embryonic trunk cells over the *HoxD* locus (mm10: chr2:74650810- 74767377). The orientations of the CBSs are as in Fig. S1. **Bottom.** Quantification of MAnorm corrected ChIP-M coverage of RAD21at the various CBS in all replicates.

**Supplemental Figure S6.**
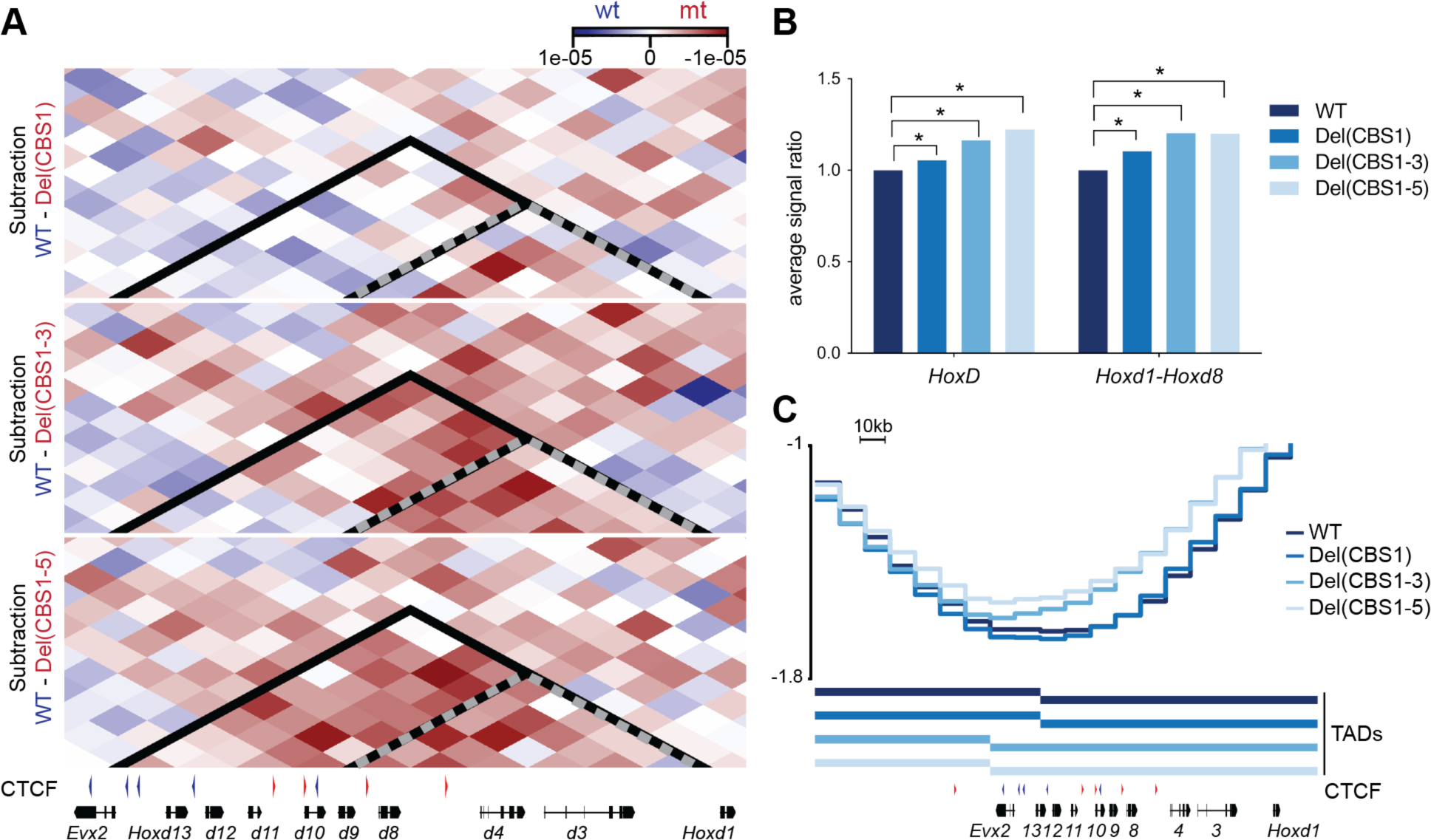
Shifting the TAD border in CBS mutants. **(A)** Subtraction maps between the observed over expected Capture Hi-C profiles from control (WT) *versus* Del(CBS1), Del(CBS1-3) and Del(CBS1-5) homozygous embryos at the *HoxD* locus (mm10: chr2:74650810-74767377). Bin size is 10kb. Blue bins represent chromatin interaction that are more prevalent in control cells whereas red bins represent interaction enriched in mutant tissues. (**B)** Bar plot showing the quantification of the interactions (ratio in comparison to the WT) within either the *HoxD* cluster (left, mm10: chr2:74660000-74760000) or the *Hoxd1-Hoxd8* region (right, mm10: chr2:74700000-74760000) in control and various mutant alleles. The asterisks represent significant changes (Mann–Whitney U test, p< 0.05). (**C)** Blue lines show the TAD separation scores in the different alleles using a window size of 240 kb. The bars below identify TADs. A shift in the position of the TAD border towards the centromeric direction is observed in the Del(CBS1-3) and Del(CBS1-5) alleles.

**Supplemental Figure S7.**
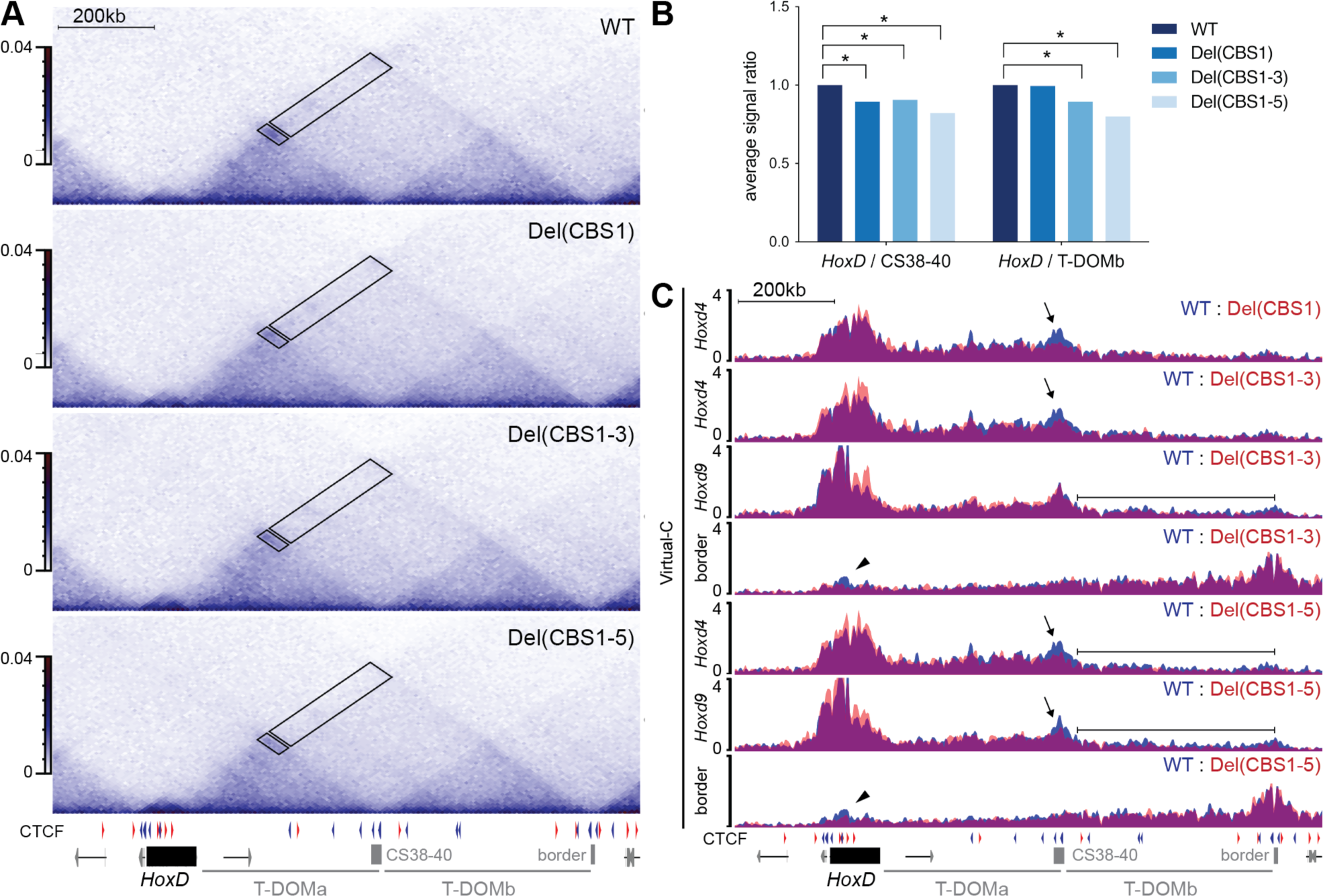
Loss of Chromatin interactions between the *HoxD* cluster and T-DOM in CBS mutants. **(A)** Capture Hi-C heatmap using E9.5 trunk cells showing the *HoxD* cluster and T-DOM (mm10: chr2:74469563-75691603) in control (WT), Del(CBS1), Del(CBS1-3) and Del(CBS1-5) samples. The orientations of the CBSs are as in Fig. S1. The position of the *HoxD* cluster (black yabox), neighboring genes (grey), the CS38-40 region and the telomeric border of T-DOMb 3’ (border) are shown below. Black boxes inside the cHi-C maps highlight the regions used for the quantifications in (**B**); the small box indicates the region used to quantify the interactions between the *HoxD* cluster (mm10: chr2:74670000-74760000) and the CS38-40 region (mm10: chr2:75120000-75160000) whereas the large box delineates the region used to quantify the interactions between the gene cluster and the T-DOMb (mm10: chr2:75170000-75590000). (**B)** Bar plot showing the ratio of interactions between the *HoxD* cluster and the CS38-40 region (left) or the T-DOMb (right) in various mutant alleles in comparison to control (WT). The asterisks represent significant changes (Mann–Whitney U test, p< 0.05). (**C)** Virtual-C with viewpoints on *Hoxd4*, *Hoxd9*, and the T-DOMb telomeric border region in control (blue) and various CBS mutant alleles (red) (mm10: chr2:74469563-75691603). The arrows indicate the loss of interactions with CS38-40 region in the Del(CBS1), Del(CBS1-3) and Del(CBS1-5) alleles. The black lines highlight the global loss of interactions with T-DOMb in the Del(CBS1-3) and Del(CBS1-5) alleles. The arrowheads point to the loss of interactions between the T-DOMb telomeric border region (border) and the gene cluster.

**Supplemental Figure S8.**
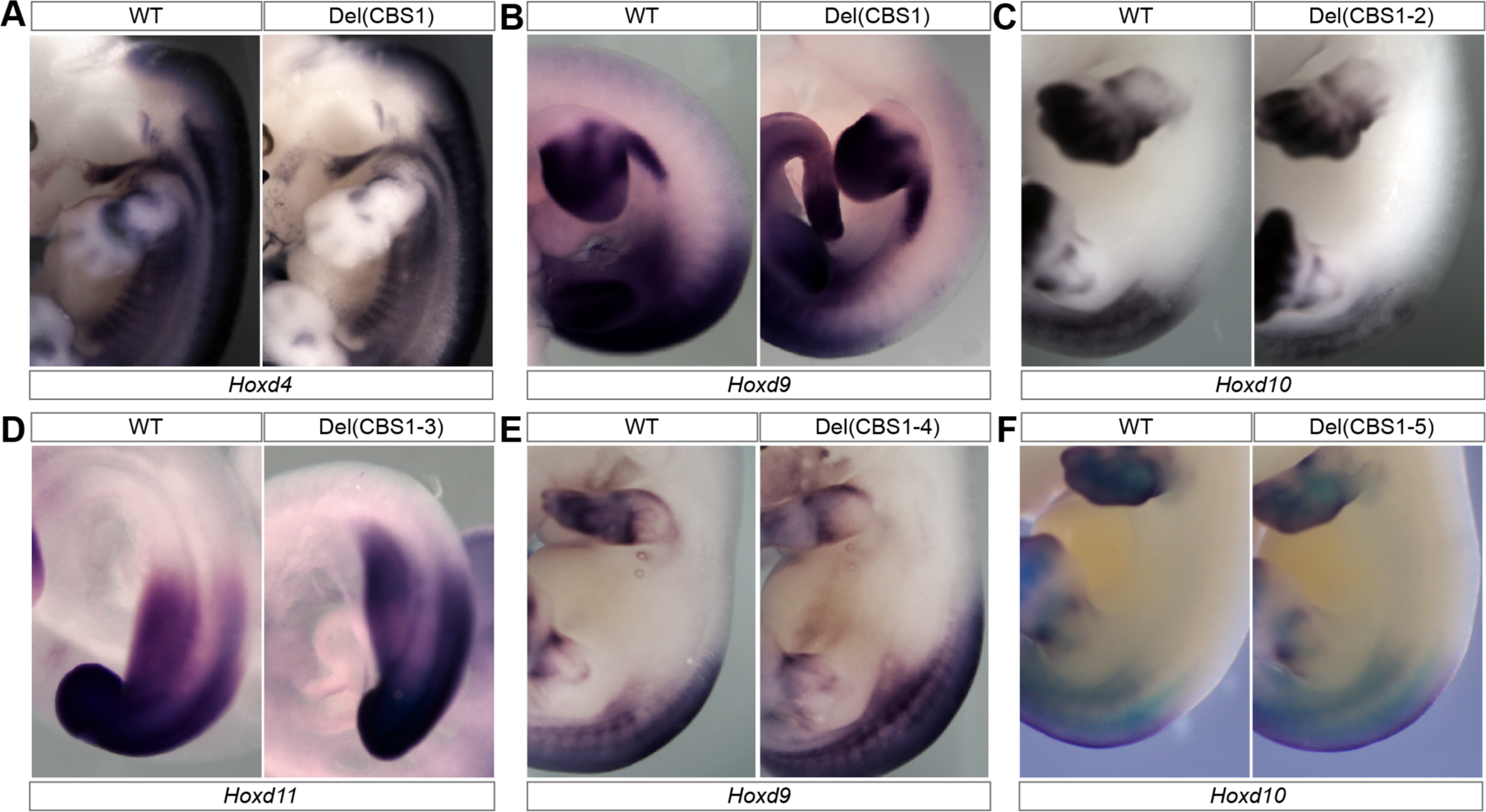
Changes in *Hoxd* genes domain of expression upon deletion of CBS. WISH showing *Hoxd* gene expression in E9.5 (D), E10.5 (B) or E12.5 (all other panels) using control (WT) embryos or various CBS homozygous mutants as indicated on top. The detected RNAs correspond to genes indicated below.

**Supplemental Figure S9.**
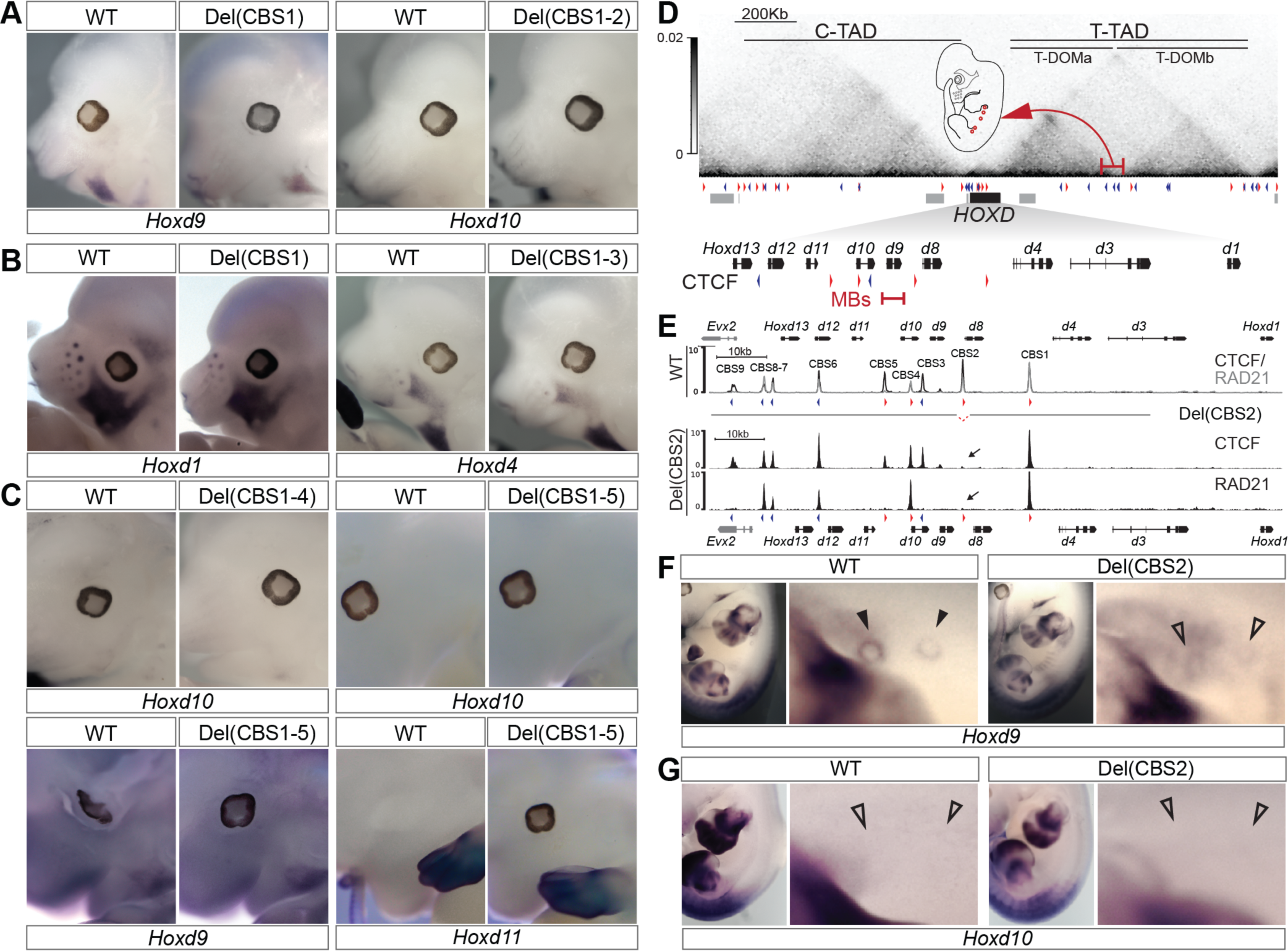
Disruption of CBS impacts *Hoxd* gene expression in vibrissae follicles, facial muscle progenitors and embryonic mammary glands. **(A-C)** WISH showing the presence of *Hoxd* mRNAs in the VFs and FMPs of either control (WT), or different CBS mutant alleles, as indicated in each panel. **(D)** Schematic representation of *Hoxd* gene regulation in the embryonic mammary gland mesenchyme (Schep et al., 2016) showing the *HoxD* locus with the position of the enhancer(s) (red bar and arrow). The enlargement of the *HoxD* cluster below highlights that only *Hoxd9* is normally expressed in these placodes. **(E) Top**. ChIP-M profiles of CTCF (black) and RAD21 (grey) over the *HoxD* cluster in control E10.5 trunk cells (as in Fig. 2). The CBS2 deletion is represented by a dashed red line below the top CTCF and RAD21 ChIP-M tracks. **Bottom**. ChIP-M tracks of CTCF (top) and RAD21 (bottom) of E10.5 Del(CBS2) mutant trunk cells (mm10: chr2:74650810-74767377). Black arrows indicate the loss of CTCF and RAD21 at CBS2 of homozygous Del(CBS2) mutant embryos. **(F-G)** WISH showing *Hoxd9 and Hoxd10* RNAs in embryonic mammary buds of either control (WT), or Del(CBS2) littermates, as indicated in each panel. Filled or empty black arrowheads indicate presence or absence, respectively, of *Hoxd9*/*Hoxd10* expression in the anterior embryonic mammary buds.

**Supplemental Figure S10.**
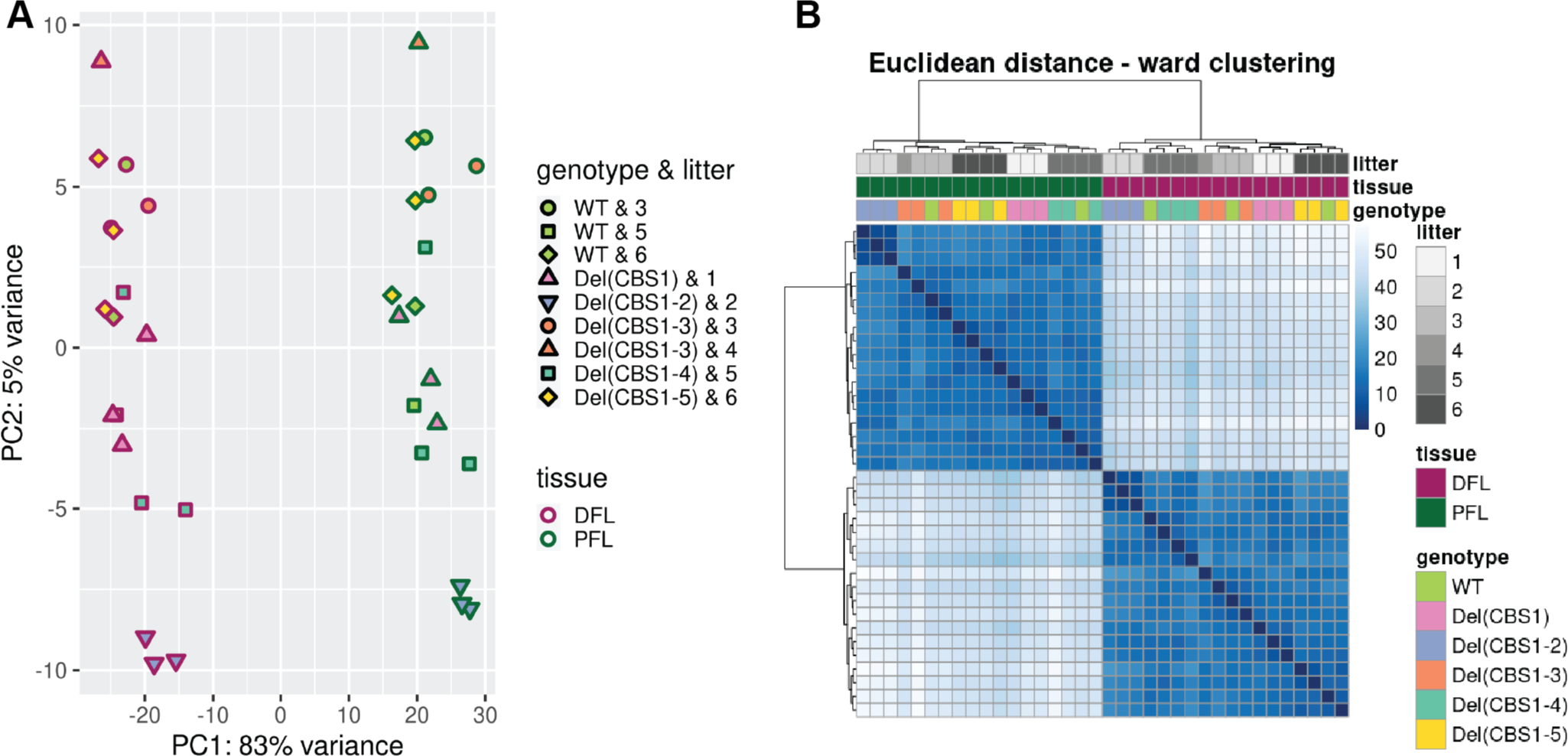
Overview of gene expression in control and CBS mutant alleles. **(A)** Principal component analysis (PCA) of gene expression levels (computed on log2(FPKM +1) of the 500 most variant autosomal protein coding genes). The genotypes and litter numbers are indicated by a combination of color fill and shape respectively (upper right corner). Tissues are indicated by outer color, with distal forelimb (DFL, pink) and proximal forelimb (PFL, green). The numbers indicate the proportion of the variance explained by PC1 or by PC2. **(B)** Hierarchical clustering (ward method) and heatmap of pairwise Euclidean distances between samples, computed on log2(FPKM +1) expression levels of the 500 most variant autosomal protein coding genes. The distances are color-coded, with dark blue representing small distances and light blue large distances. Litter numbers, tissues and genotypes are color coded as indicated in the right side of the image.

**Supplemental Figure S11.**
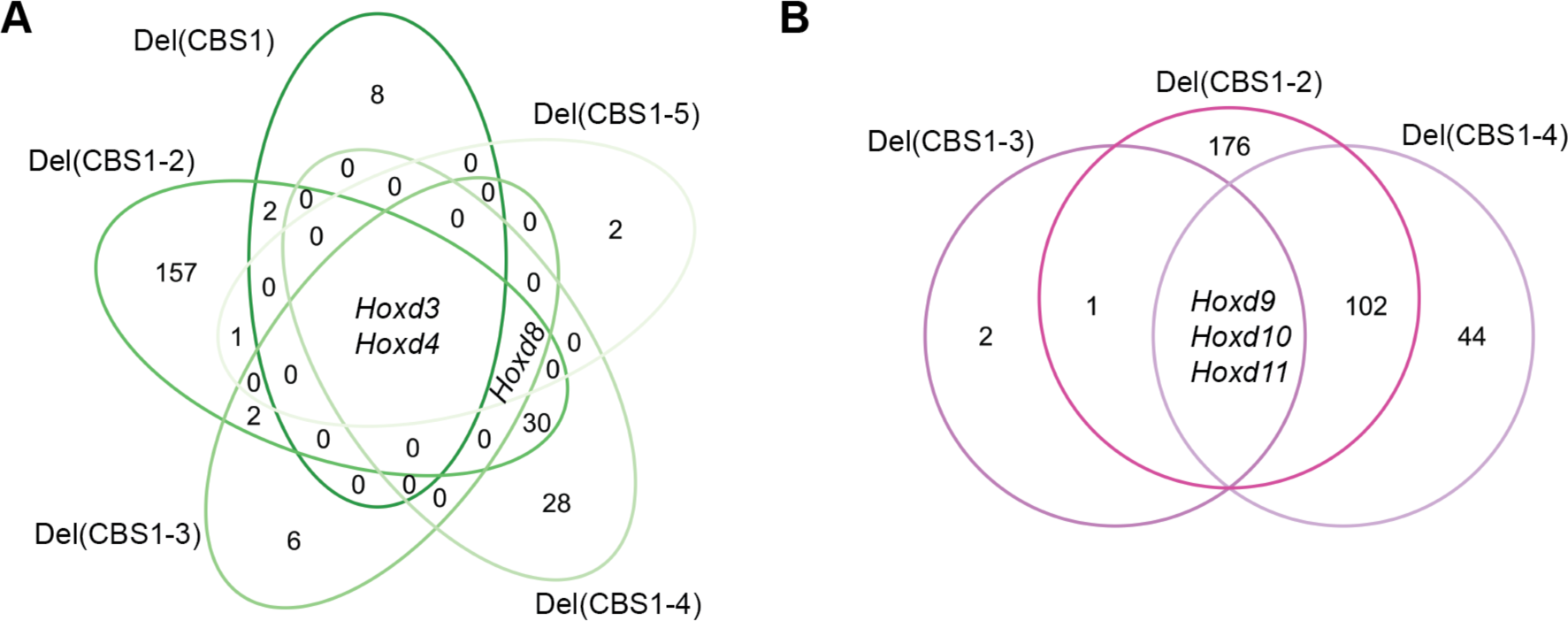
Differential gene expression amongst CBS mutants. **(A-B)** Venn diagrams of pairwise differential expression analyses of autosomal protein coding genes (absolute log2 fold change > 0.58 and adjusted p-value < 0.05) for proximal forelimb cells **(A)** and distal forelimb cells **(B)** in selected CBS mutant alleles. *Hoxd3* and *Hoxd4* are the only commonly differentially expressed genes amongst all CBS alleles in proximal forelimb cells **(A)** and *Hoxd8* the only one shared by four out of five CBS alleles. *Hoxd9*, *Hoxd10*, and *Hoxd11* are the only commonly differentially expressed genes between the Del(CBS1- 2), Del(CBS1-3) and Del(CBS1-4) alleles in DFL **(B)**.

**Supplemental Figure S12.**
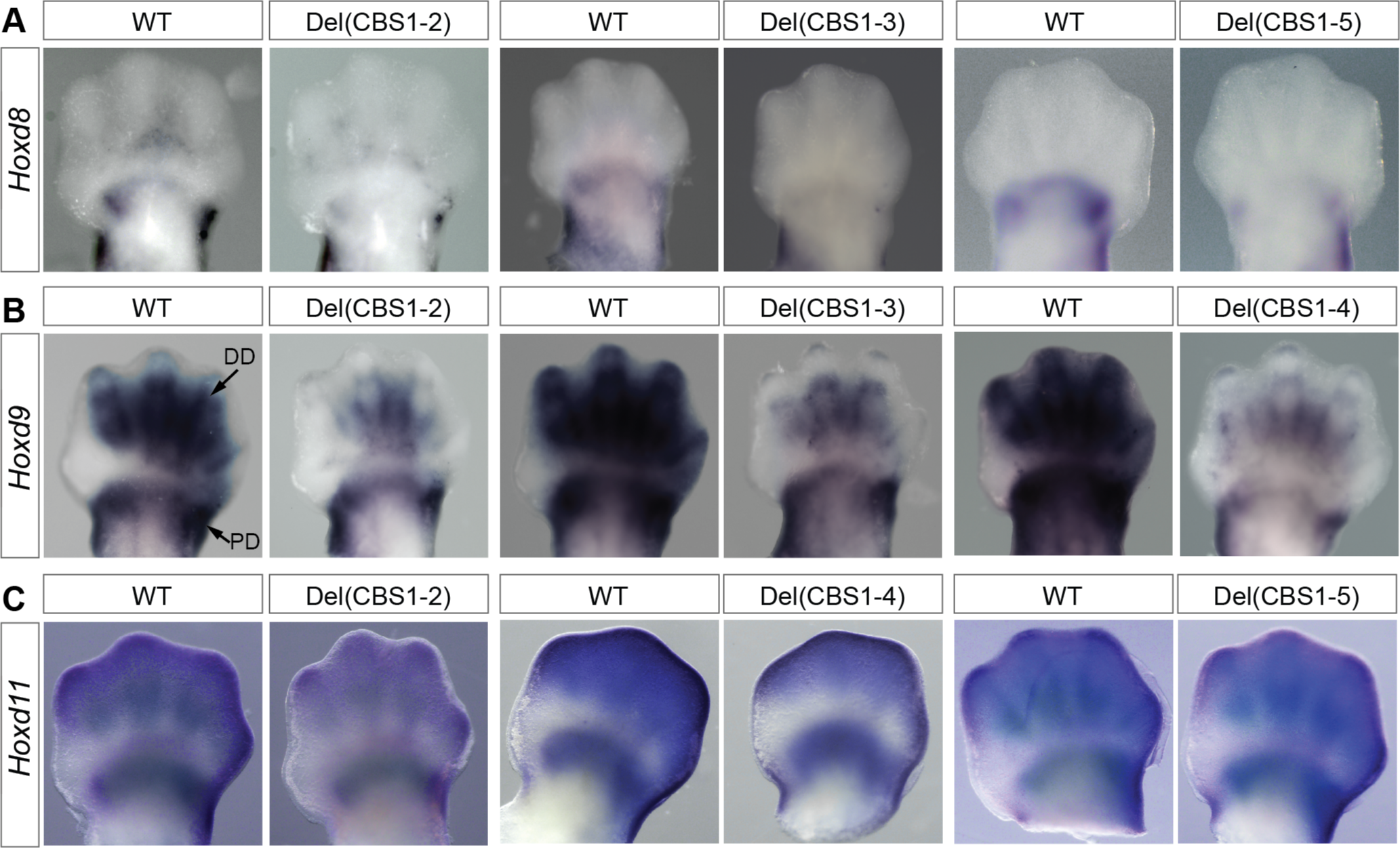
Disruption of CBS impacts *Hoxd* gene expression developing limb buds. **(A-C)** WISH showing *Hoxd* gene expression in E12.5 forelimbs of control (WT) or various CBSs mutant embryos, as indicated in each panel. The genes analyzed are indicated on the left. The proximal (PD) and distal (DD) expression domains are indicated by using the control *Hoxd9* panel in **(B)** (left).

**Supplemental Figure S13.**
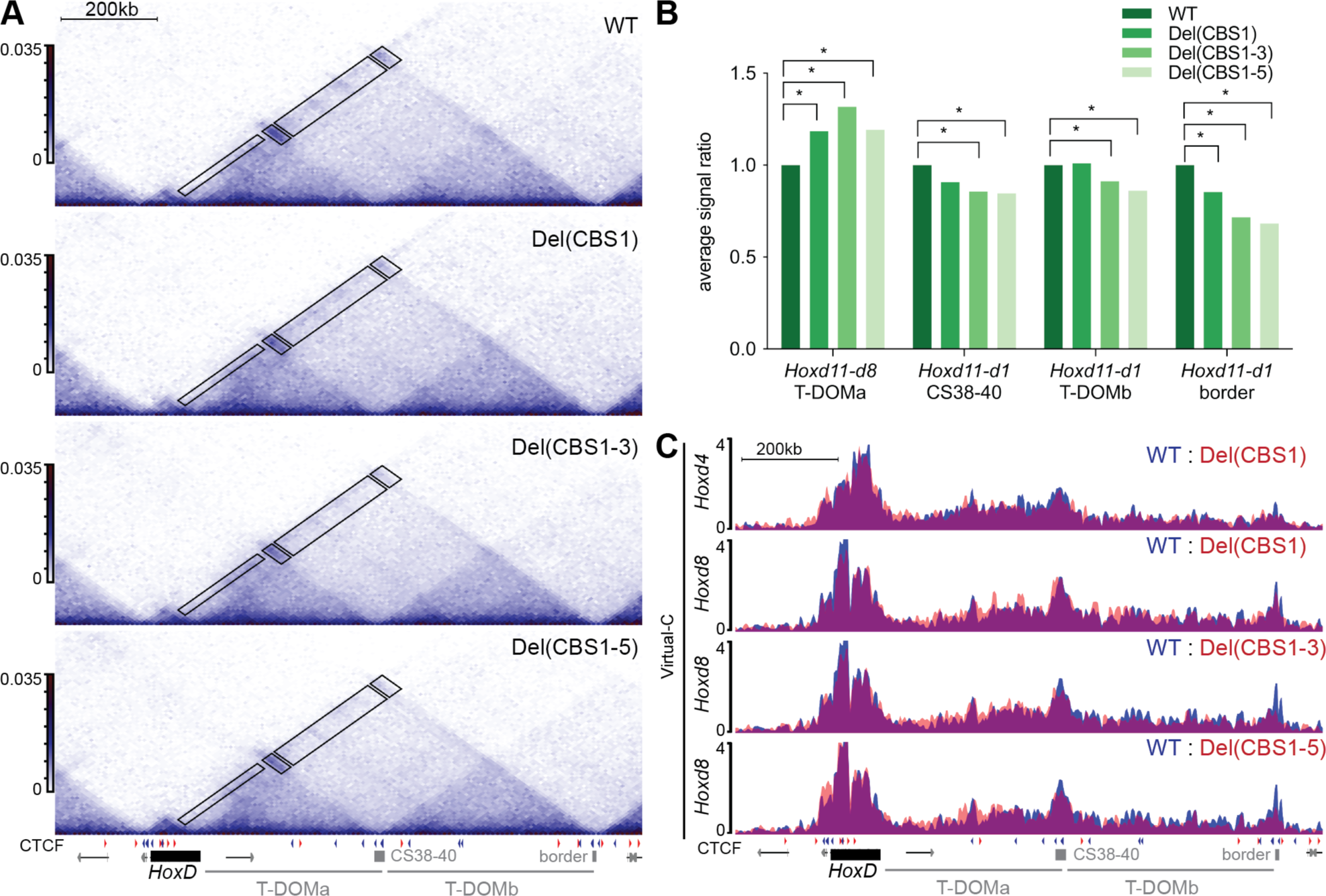
Differences in chromatin structure between control and CBS mutant proximal forelimb cells. **(A)** Capture Hi-C map of E12.5 proximal forelimb buds showing the *HoxD* cluster and T-DOM (mm10: chr2:74469563-75691603) in control (WT), Del(CBS1), Del(CBS1-3) and Del(CBS1-5) mutant specimen. CTCF sites and their orientations are shown below as in Fig. S1. Black boxes on the cHi-C maps highlight the regions used for the quantifications shown in **(B)**. **(B**) From left to right, *Hoxd11-d8*/T-DOMa, *Hoxd11-d1*/CS38-40, *Hoxd11-d1*/T-DOMb, and *Hoxd11-1*/T-DOMb 3’ border. Bar plot displaying the changes in interaction ratio, when compared to control, between the *HoxD* cluster and the regions boxed in **(A)**, using cells from in WT, Del(CBS1), Del(CBS1-3) and Del(CBS1-5) mutant alleles. Coordinates for the regions are (mm10): *Hoxd11 to Hoxd8*: chr2:74680000-74710000, T- DOMa: chr2:74770000-75100000, *Hoxd11 to Hoxd1*: chr2:74680000-74760000, CS38-40: chr2:75120000-75160000, T-DOMb: chr2:75170000-75560000, T-DOMb 3’ border: chr2:75570000- 75620000. The asterisks represent significant changes (Mann–Whitney U test, p< 0.05). **(C)** Virtual-C with viewpoints selected on *Hoxd4* and *Hoxd8* using control (WT, blue), Del(CBS1), Del(CBS1-3) and Del(CBS1-5) alleles (red) (mm10: chr2:74469563-75691603).

**Supplemental Table S1.** Incidence of vertebral column defects in mice carrying *in-cis* mutations in CTCF sites within the *HoxD* cluster.

|  | Occipital-C1 ventral malformations<sup>1</sup> | C1 or 2 dorsal malformations<sup>2</sup> | C1 or 2 ventral malformations<sup>3</sup> | C6-T1 transverse malformations<sup>4</sup> | L5 transverse malformations<sup>5</sup> |
| --- | --- | --- | --- | --- | --- |
| WT | 0/6 | 0/6 | 0/6 | 0/6 | 0/6 |
| Del(CBS1-5) | 7/9 | 7/9 | 6/9 | 0/9 | 1/9 |
| Del(CBS1-4) | 6/10 | 5/10 | 5/10 | 2/11 | 4/11 |
| Del(CBS1-3) | 4/5 | 4/5 | 4/5 | 3/5 | 0/5 |

<sup>1</sup> Variable combinations of basi-occipital bone and atlas anterior tubercle and/or anterior arch malformations, fusions, or supernumerary ossified elements.
<sup>2</sup> Variable combinations of atlas and axis neural arch overgrowth, fusions or supernumerary ossified elements.
<sup>3</sup> Variable combinations of atlas anterior arch and axis odontoid process fusions.
<sup>4</sup> Variable combinations of C6, C7 cervical ribs associated to next posterior C7 or T1 ribs respectively.
<sup>5</sup> Presence of only four normal lumbar vertebrae, the fifth and or sixths showing asymmetrical or complete transformation to the sacral form.

**Supplemental Table S2.**
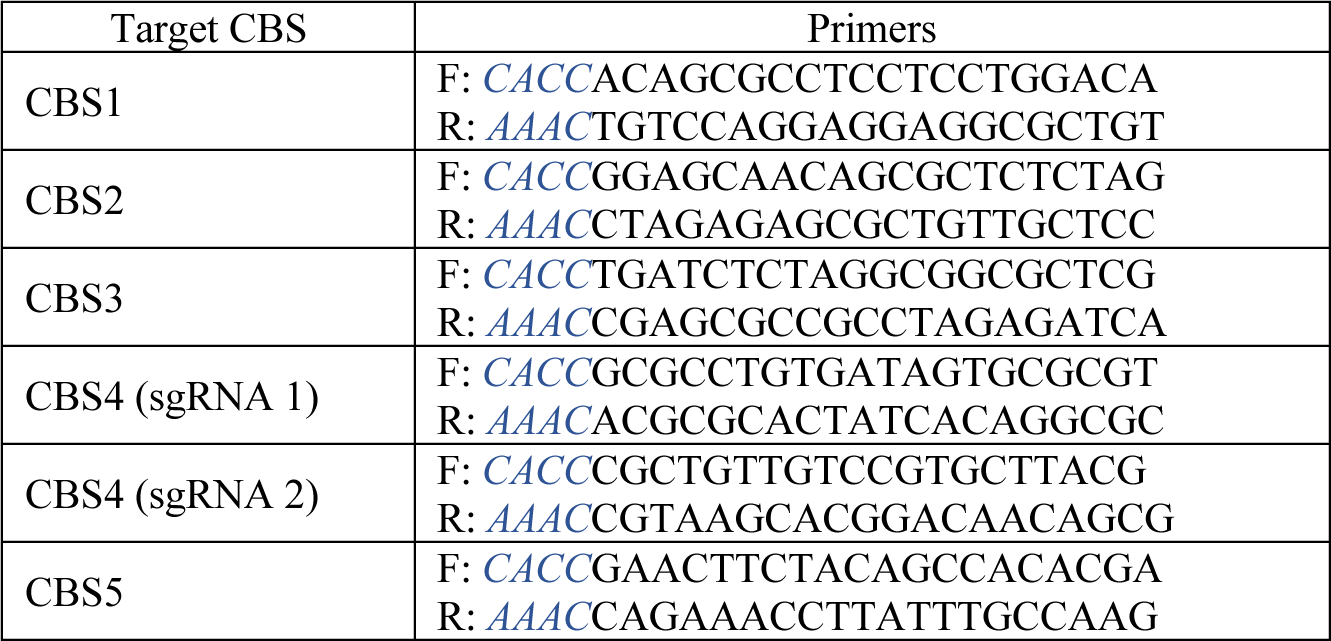
List of sgRNAs used to target the various CBSs.

**Supplemental Table S3.**
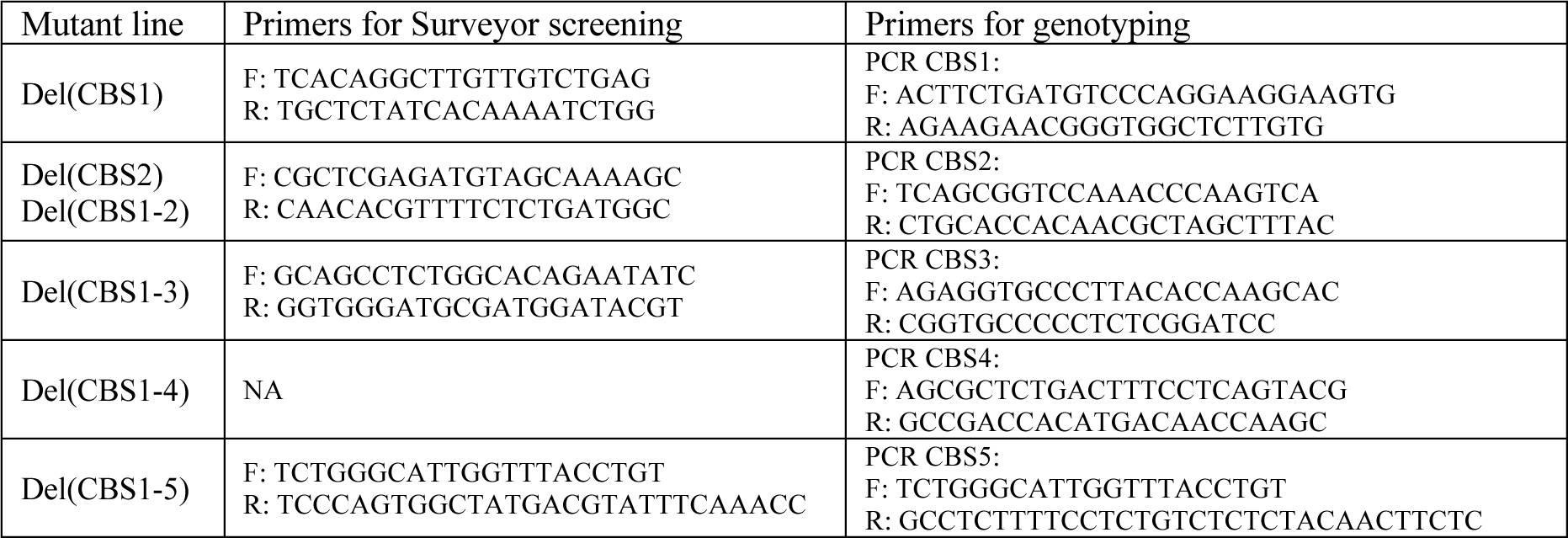
List of primers used for characterizing and genotyping the mutations induced at various CBSs.

**Supplemental File 1**: Summary of pairwise differential gene expression. Table reporting for each gene in which pairwise differential gene expression analysis the gene was significant. For each analysis where it was significant, the associated log2 fold-change and adjusted p-value are provided.

